# Entry by multiple picornaviruses is dependent on a pathway that includes TNK2, WASL and NCK1

**DOI:** 10.1101/737635

**Authors:** Hongbing Jiang, Christian Leung, Stephen Tahan, David Wang

**Author notes:** Address correspondence to: Hongbing Jiang and David Wang, Washington University in St. Louis School of Medicine, Campus box 8230, 660 S. Euclid Ave., St. Louis, MO 63110 USA.

## Abstract

Comprehensive knowledge of the host factors required for picornavirus infection would facilitate antiviral development. Here we demonstrate roles for three human genes, TNK2, WASL, and NCK1, in infection by multiple picornaviruses. CRISPR deletion of TNK2, WASL or NCK1 reduced encephalomyocarditis virus (EMCV), coxsackievirus B3 (CVB3), poliovirus and enterovirus D68 infection, and chemical inhibitors of TNK2 and WASL decreased EMCV infection. Reduced EMCV lethality was observed in mice lacking TNK2. TNK2, WASL and NCK1 were important in early stages of the viral lifecycle, and genetic epistasis analysis demonstrated that the three genes function in a common pathway. Mechanistically, reduced internalization of EMCV was observed in TNK2 deficient cells demonstrating that TNK2 functions in EMCV entry. Domain analysis of WASL demonstrated that its actin nucleation activity was necessary to facilitate viral infection. Together, these data support a model wherein TNK2, WASL, and NCK1 comprise a pathway critical for multiple picornaviruses.

## Introduction

Picornaviruses cause a wide range of diseases including the common cold, hepatitis, myocarditis, poliomyelitis, meningitis, and encephalitis (1). Although vaccines exist for poliovirus and hepatitis A virus, there are currently no FDA approved antivirals against picornaviruses in the United States. As obligate intracellular pathogens, viruses are dependent on the host cellular machinery to complete their lifecycle. Targeting of such host cellular factors in the design of antiviral drugs can circumvent resistance that arises from rapid mutation of viruses. The early stages of the virus lifecycle, including receptor binding, entry, uncoating, and initiation of replication, are the ideal targets for preventing virus infection since the virus has not yet multiplied.

Despite extensive studies (2), there remain significant gaps in our understanding of virus entry. As the family *Picornaviridae* encompasses a wide range of viruses, it is not surprising that there is diversity in the known entry mechanisms of different species. Among the picornaviruses, poliovirus entry has been the most extensively studied. While some reports suggest that poliovirus enters the cell through clathrin-mediated endocytosis and that its genome release depends on endosome acidification (3), more recent studies report that poliovirus enters cells by a clathrin-, caveolin-, flotillin-, and microtubule-independent pathway (4). Furthermore, poliovirus entry is sensitive to inhibitors of both tyrosine kinases and actin-polymerization, although it is not known which specific tyrosine kinase(s) is/are critical for poliovirus infection (4). Coxsackie virus B3 (CVB3) entry has also been extensively studied (5). In polarized epithelial cells, CVB3 binding to the co-receptor decay-accelerating factor (DAF) and the coxsackievirus and adenovirus receptor (CAR) leads to entry by caveolin-dependent endocytosis and macropinocytosis (6, 7). In contrast to CVB3 and poliovirus, there have been few studies of EMCV entry. Vascular cell adhesion molecule 1 (VCAM-1) is reported to be a receptor for EMCV (8). Interaction of the EMCV virion with VCAM-1 is believed to induce a conformational change that then releases the viral RNA genome; entry into the cytosol is reported to be independent of acidification (9).

Using a novel virus infection system comprised of the model organism *C. elegans* and Orsay virus, the only known natural virus of *C. elegans*, we previously identified several genes that are essential for virus infection in *C. elegans* (10). The genes *sid-3, viro-2* and *nck-1* were found to be essential for an early, pre-replication step of the Orsay virus lifecycle. *sid-3* encodes a non-receptor tyrosine kinase orthologous to human Tyrosine Kinase Non-Receptor 2 (TNK2), *viro-2* encodes an orthologue of human Wiskott-Aldrich Syndrome protein Like protein (WASL), and *nck-1* encodes an orthologue of Non-Catalytic Region of Tyrosine Kinase (NCK1), an adaptor protein that binds to both TNK2 and WASL (11, 12). Since Orsay virus is a non-enveloped, positive strand RNA virus that is evolutionarily related to the family *Picornaviridae*, we reasoned that the human orthologues of these genes may have a conserved role in infection by picornaviruses.

Human TNK2 is linked to cancers, has been reported to be activated by multiple extracellular stimuli, and is involved in many different pathways, such as clathrin/receptor mediated endocytosis, regulation of EGFR degradation, transduction of signals into the nucleus, and regulation of actin polymerization (11, 13-15). Furthermore, overexpression of TNK2 induces interferon and leads to reduced replication of a Hepatitis C virus replicon (16). To date, there are no publications demonstrating a positive role for TNK2 in virus infection; several siRNA based screens suggest that TNK2 is important for influenza A virus (IAV), vesicular stomatitis virus (VSV), and hepatitis C virus (HCV) infections (16-19), but no validation of these screening results have been reported. Humans encode two orthologues of *viro-2*, WASP and WASL (Wiskott-Aldrich syndrome protein/-like) (20). Strikingly, biochemical assays have demonstrated that WASL is a substrate for the kinase activity of TNK2 (15), suggesting that the two function in a pathway. NCK1 is an adaptor protein reported to co-localize with TNK2 (11, 21) and interact with WASL. There are only limited studies linking WASL and NCK1 to virus infection. A clear role has been established for WASL in spread of vaccinia virus (22) that involves NCK1 as well (12, 23). Furthermore, sensitivity of Lassa virus infection to Wiskostatin, a small molecule inhibitor of WASL, suggests a role of WASL in virus entry (24). However, there are no studies implicating NCK1 in any aspect of picornavirus infection or in entry of any virus.

Here, we determined whether these three human orthologues of the genes found in a *C. elegans* forward genetic screen function in an evolutionarily conserved manner to facilitate virus infection in human cell culture. CRISPR-Cas9 genome editing was used to generate knockout cells for each gene, and their impact on infection by a panel of viruses from different families was then tested. Significant reductions in infection by multiple picornaviruses were observed in each of the cell lines. Consistent with these *in vitro* data, we observed increased survival of mice lacking murine TNK2 after EMCV challenge *in vivo*. We further demonstrated that TNK2, WASL and NCK1 function in a pathway to support EMCV virus infection with WASL and NCK1 lying downstream of TNK2. Mechanistically, loss of TNK2 led to reduced EMCV virus internalization, while both TNK2 and WASL were required for proper endocytic trafficking of EMCV. These data support a model wherein TNK2, WASL and NCK1 comprise a pathway that is critical for entry by multiple picornaviruses.

## Results

### Non-receptor tyrosine kinase TNK2 is critical for infection by multiple picornaviruses

To investigate the function of TNK2 in virus infection, human lung epithelial carcinoma A549 cells deficient in TNK2 were generated by CRISPR-Cas9 genome editing (Figure 1A). We tested a panel of viruses including multiple viruses from the family *Picornaviridae*, which are evolutionarily related to Orsay virus (25). In a single step growth analysis, deletion of TNK2 by two independent sgRNAs (TNK2 KO1 and TNK2 KO2 cells) reduced EMCV infection by 93% and 86%, respectively, compared to control cells (P<0.00001, Figure 1B). When infected by CVB3, 56% and 48% fewer infected TNK2 KO1 and TNK2 KO2 cells were observed (P<0.0001, Figure 1C). In a multi-step growth curve, EMCV titers in the supernatant from TNK2 KO1 were 1660-fold lower at 24 hours post infection (P<0.01, Figure 1D). In addition, reduced infectivity in TNK2 KO1 was also observed with both recombinant GFP-EMCV and GFP-CVB3, respectively (Figure 1-figure supplement1 A and B). Furthermore, we examined virus replication complex formation by double stranded RNA immunostaining with the commercial J2 antibody and direct electron microscopy (EM) on EMCV infected cells. We observed fewer cells positive for double stranded RNA and smaller size of the virus replication complex (Figure 1-figure supplement1 C and D). To corroborate these findings, we analyzed EMCV and CVB3 infection in commercial Hap1 cells (a haploid hematopoietic derived lymphoma cell line) deficient in TNK2. Reductions in infected cells by 24% for EMCV and 30% for CVB3 were observed (P<0.001, Figure1-figure supplement2 A and B). Statistically significant reductions in poliovirus infection in both Hap1 and A549 TNK2 knockout cells and reduction in enterovirus D68 infection in A549 cells were also observed (Figure 1-figure supplement2 C, E and F). In contrast, no effect of TNK2 deletion was seen for IAV, parainfluenza virus (PIV5) or adenovirus 5 (Figure 1-figure supplement2 D, G, H and I), demonstrating the apparent specificity of TNK2 for picornavirus infection.

**Figure 1.**
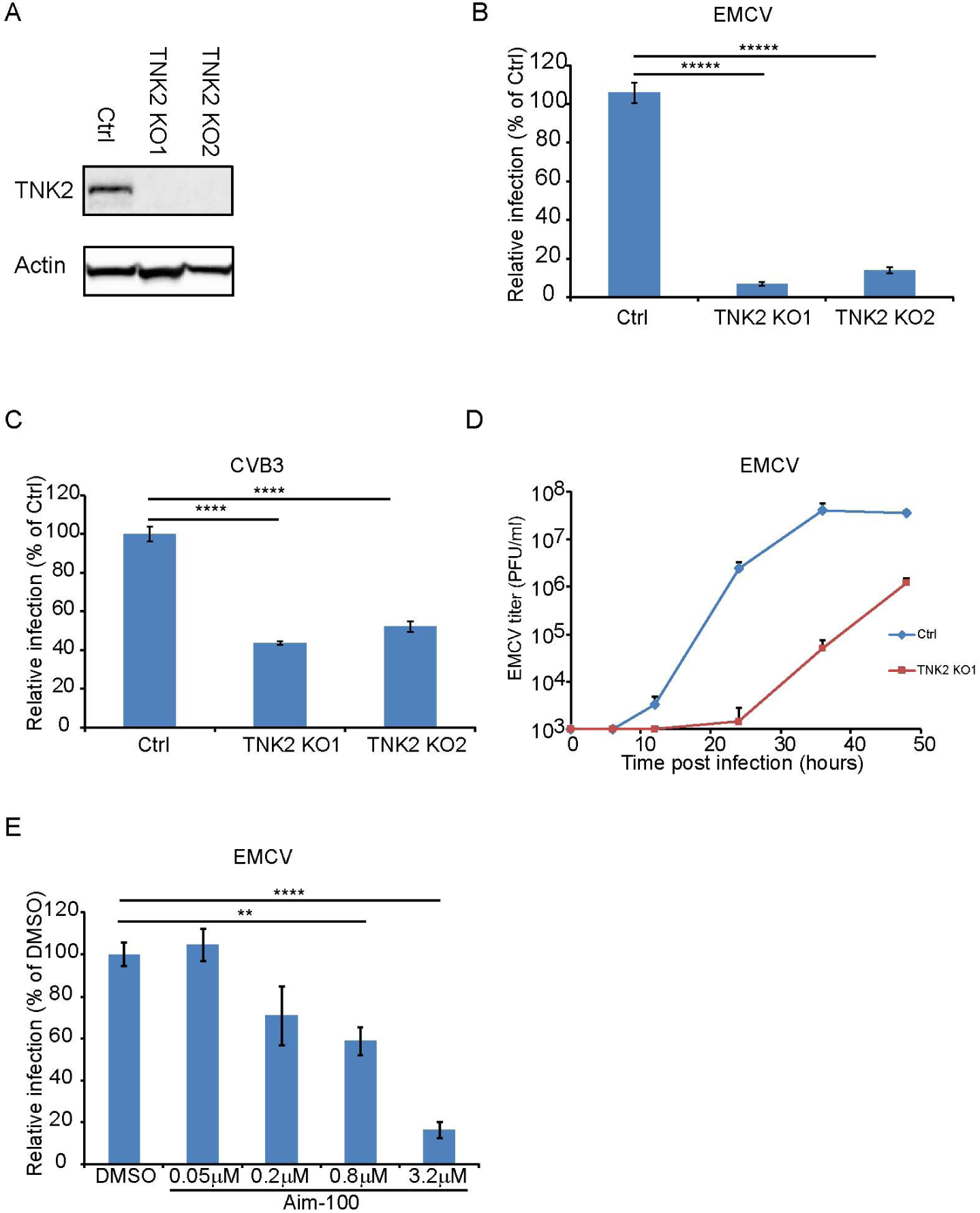
TNK2 is critical for multiple picornavirus infections. (A) TNK2 protein expression in TNK2 KO1, TNK2 KO2 and Ctrl (control) cells generated by CRISPR-Cas9 genome editing with either specific targeting or non-specific targeting sgRNA in A549 cells. Cells lysates were analyzed by Western blot. (B) FACS quantification of EMCV positive cells for TNK2 KO1, TNK2 KO2, and Ctrl cells 10 hours post infection at an MOI of 1. (C) FACS quantification of CVB3 virus positive cells for TNK2 KO1, TNK2 KO2, and Ctrl cells 8 hours post infection at an MOI of 1. (D) Multi-step growth curve for EMCV multiplication on TNK2 KO1 and Ctrl cells at an MOI of 0.01. Virus titers in the culture super-natant were quantified by plaque assay at 0, 6, 12, 24, 36, and 48 hours post infection. (E) Aim-100 inhibition of EMCV infection on naïve A549 cells. A549 cells were pre-treated with Aim-100 at indicated concentrations and infected with EMCV at an MOI of 1. Virus positive cells were quantified by FACS. (B-E) Error bars represent standard deviation of three replicates. The data shown are representatives of three independent experiments. *: P<0.05, ***: P<0.001, ****: P<0.0001, *****: P<0.00001.

We next attempted to rescue the EMCV infection defect in TNK2 knockout cells by ectopic overexpression of TNK2. However, the complexity of the TNK2 locus (three major isoforms and 22 putative transcript variants (26)) presented challenges, and the reported induction of interferon by TNK2 overexpression (16) may have further complicated interpretation of these experiments. Lentivirus expression of each of the three major TNK2 isoforms (which yielded expression levels of TNK2 much higher than the endogenous levels (Figure 1-figure supplement3 D)) in control A549 cells led to statistically significant reductions in EMCV infection (Figure 1-figure supplement3 A, B and C). In addition, fluorescently tagged TNK2 expression through either direct transfection in 293T cells or lentivirus transduction in A549 cells led to aggregates inside the cytosol (Figure 3-figure supplement2 and Figure 5-figure supplement A). Nevertheless, we consistently observed a small, but statistically significant increase in EMCV virus infection in TNK2 KO1 cells transduced with the canonical isoform 1 as compared to the control cells (Figure 1-figure supplement3 A). We also used an alternative approach to rescue virus infection in TNK2 KO1 cells by reverting the 28 bp deletion in clone TNK2 KO1 using CRISPR-Cas9 with oligonucleotide mediated homologous template directed recombination (HDR); 3 synonymous mutations were included in the template to unambiguously identify a successful repair event (Figure 1-figure supplement3 H). Screening of 500 single-cell clones yielded one clone that had repaired the TNK2 28 bp deletion allele. However, this process also introduced an insertion of 11 nucleotides in the adjacent intron sequence 9 bp from the nearby splice acceptor (Figure 1-figure supplement3 H). We were able to restore low levels of TNK2 expression in the HDR repaired clone (Figure 1-figure supplement3 G). In this clone, EMCV infected cell percentage and virus titer increased, by 8-fold (P<0.0001, Figure 1-figure supplement3 E) and 9-fold (P<0.01, Figure 1-figure supplement3 F), respectively as compared to TNK2 KO1 cells.

As a completely independent means of assessing the role of TNK2, we treated A549 cells with Aim-100, a small molecule kinase inhibitor with specificity for TNK2 (27). Aim-100 pre-treatment reduced EMCV infection in a dose dependent manner without affecting cell viability (Figure 1E and data not shown).

### WASL, a known substrate of TNK2, is critical for picornavirus infection

The human genome encodes two WASP paralogues, WASP and WASL (also known as N-WASP). In this study, we focused on WASL because WASP is not expressed in epithelial cells such as A549. Clonal WASL knockout A549 cells were generated by CRISPR-Cas9 genome editing (Figure 2A). In a single step growth analysis, WASL KO cells showed 62% reduction in EMCV infected cells as compared to control cells (P<0.0001, Figure 2B), while in a multi-step growth curve, WASL KO cells yielded 248-fold reduced virus titers at 24 hours post infection (P<0.01, Figure 2D). Ectopic expression of WASL in the WASL KO cells rescued EMCV infection to wild type levels (Figure 2-figure supplement A and B). As with TNK2, we observed fewer cells positive for double stranded RNA by immunostaining and smaller size of the virus replication complex by EM (Figure 1-figure supplement1 C and D). Reduced levels of CVB3, poliovirus and enterovirus D68 infection were observed in these cells (Figures 2C, Figure1-figure supplement2 E, F and M). In addition, independent Hap1 cells deficient in WASL had reduced EMCV, CVB3 and poliovirus infection (Figures 1-figure supplement2 A, B and C). To independently assess the role of WASL during virus infection, we used a small molecule inhibitor of WASL, Wiskostatin (28). Wiskostatin reduced EMCV infection in A549 cells in a dose dependent manner with no apparent decrease of cell viability (Figure 2E and data not shown). These data demonstrate that WASL is critical for multiple picornavirus infection in human cell culture.

**Figure 2.**
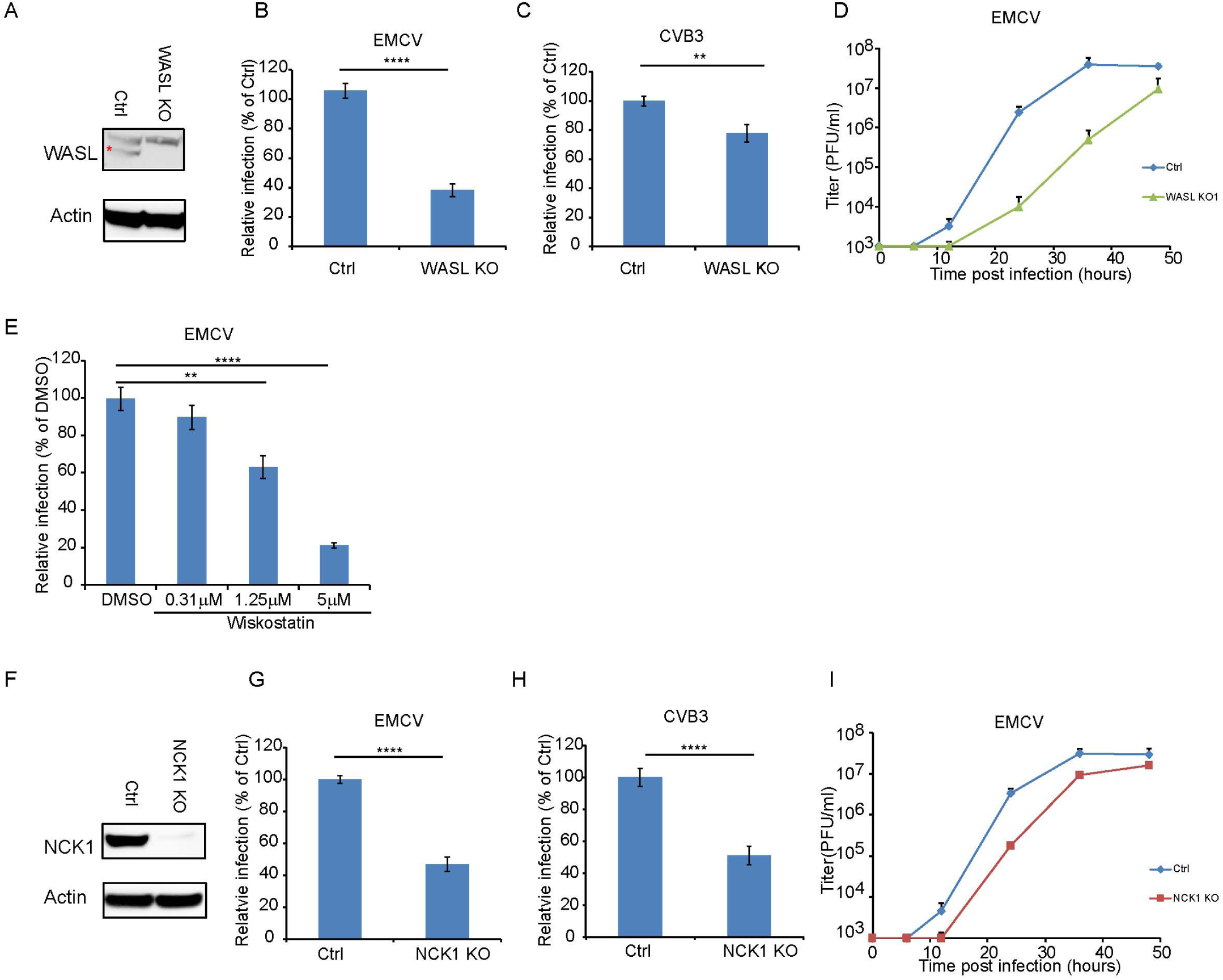
WASL and NCK1 are critical for multiple picornavirus infections. (A) WASL protein expression in WASL KO and Ctrl cells generated by CRISPR-Cas9 genome editing with either specific targeting or non-specific targeting sgRNA in A549 cells. Cells lysates were analyzed by Western blot. (B) FACS quantification of EMCV positive cells for WASL KO and Ctrl cells 10 hours post infection at an MOI of 1. (C) FACS quantification of CVB3 positive cells for WASL KO and Ctrl cells 8 hours post infection at an MOI of 1. (D) Multi-step growth curve for EMCV multiplication on WASL KO and Ctrl cells infected at an MOI of 0.01. (E) Wiskostatin inhibition of EMCV infection on naïve A549 cells. A549 cells were pre-treated with Wiskotstatin at indicated concentrations and infected with EMCV at an MOI of 1. Virus positive cells were quantified by FACS. (F) NCK1 protein expression in NCK1 KO and Ctrl cells generated by CRISPR-Cas9 genome editing with either specific targeting or non-specific targeting sgRNA in A549 cells. Cells lysates were analyzed by Western blot. (G) FACS quantification of EMCV positive cells for NCK1 KO and Ctrl cells 10 hours post infection at an MOI of 1. (H) FACS quantification of CVB3 positive cells for NCK1 KO and Ctrl cells 8 hours post infection at an MOI of 1. (I) Multi-step growth curve for EMCV multiplication on NCK1 KO and Ctrl cells infected at an MOI of 0.01. (A) The red asterisk indicates WASL protein band. (B, C, E, G, H) Error bars represent standard deviation of three replicates. The data shown is representatives of at least two independent experiments. **: P<0.01, ***: P<0.001, ****: P<0.0001, *****: P<0.00001, NS: not significant (P>0.05).

### The signaling adaptor protein NCK1 is also critical for picornavirus infection

Deletion of NCK1 in A549 cells (Figure 2F) reduced the number of EMCV and CVB3 infected cells by 55% and 49%, respectively (P<0.0001, Figure 2G and 2H) comparable to the reductions observed in WASL KO cells. By a multi-step growth analysis, NCK1 KO showed 10-fold reduction of EMCV virus titer at 24 hours post infection (P<0.01 Figure 2I). Ectopic expression of NCK1 in the NCK1 KO cells rescued EMCV infection to wild type levels (Figure 2-figure supplement C and D). As with TNK2 and WASL, these data demonstrate that NCK1 is critical for multiple picornavirus infection in human cell culture.

### TNK2, WASL, and NCK1 are components of a common pathway

To determine whether TNK2, WASL, and NCK1 act in a common or distinct pathway for virus infection, we performed genetic epistasis analysis by generating double and triple mutant cell lines (Figure 3-figure supplement1 A). EMCV infection levels and viral titer production in the double and triple knockout lines were the same as observed in the TNK2 KO1 lines (Figure 3A and 3B). The lack of an additive effect suggests that WASL and NCK1 are both in the same genetic pathway as TNK2. Combined with the known ability of TNK2 to phosphorylate WASL (15), these data support a model where TNK2 acts upstream of WASL and NCK1. The magnitude of the impact of TNK2 KO1 was greater than that of WASL KO or NCK1 KO, suggesting that TNK2 might work through additional pathways in mediating virus infection besides acting through WASL and NCK1.

**Figure 3.**
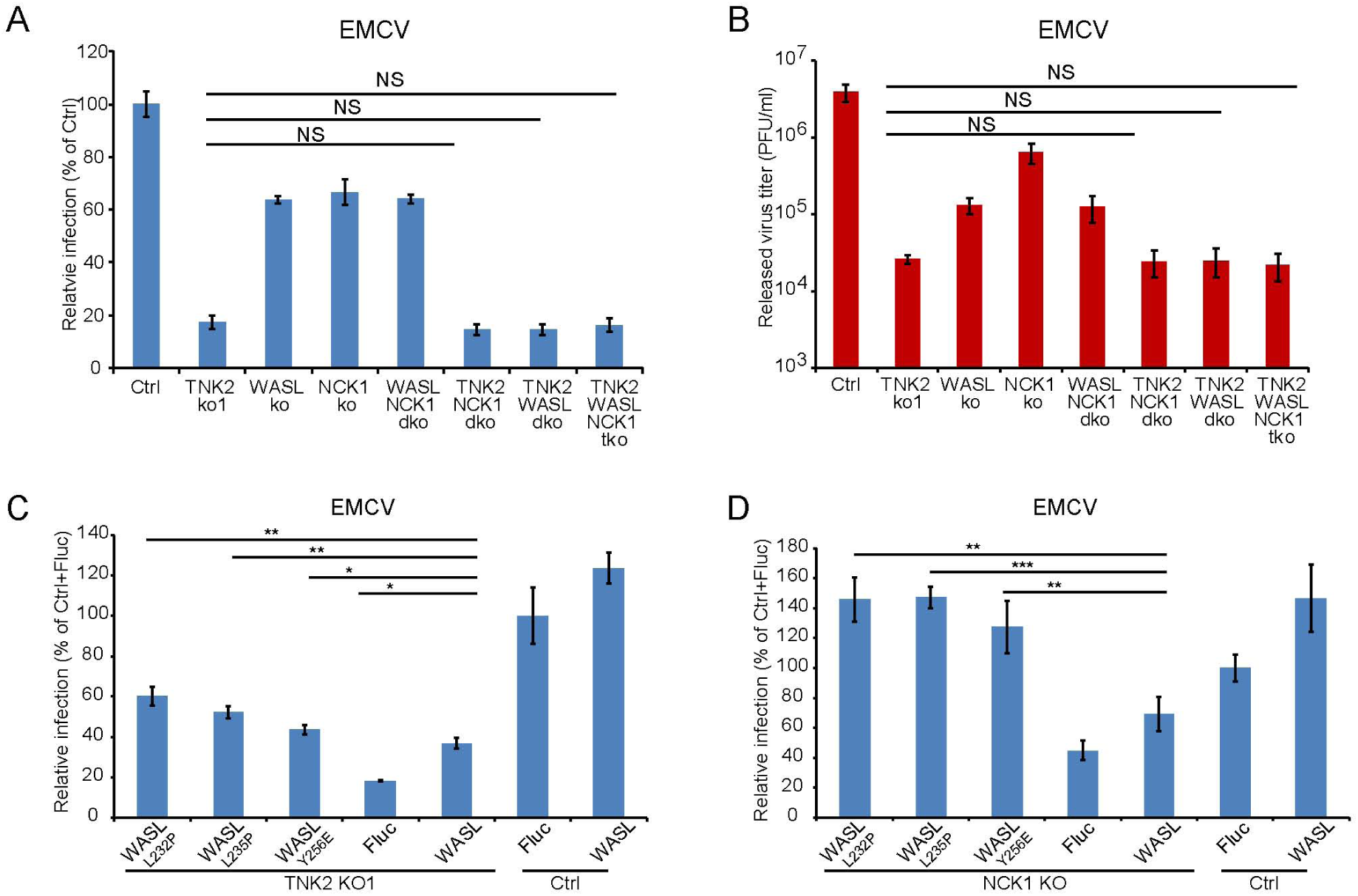
TNK2, WASL, and NCK1 are in a pathway supporting virus infection. (A) FACS quantification of EMCV positive cells for TNK2, WASL, NCK1 single, double, triple gene knockout and Ctrl cells 10 hours post infection at an MOI of 1. (B) Virus titer for EMCV multiplication on TNK2, WASL, and NCK1 single, double, triple gene knockout and Ctrl cells at 24 hours post infection at an MOI of 0.01. (C) FACS quantification of EMCV positive cells for TNK2 KO1 cells that were transduced with constitutively active WASL constructs 10 hours post infection at an MOI of 1. (D) FACS quantification of EMCV positive cells for NCK1 KO cells that were transduced with constitutively active WASL constructs 10hours post infection at an MOI of 1. (A, B) dko: double knockout, tko: triple knockout. (B-D) Error bars for virus infection represent standard deviation of three replicates. The data shown is representatives of two independent experiments.

Because TNK2 phosphorylation of WASL increases it actin nucleation activity (29), we reasoned that it might be possible to complement the TNK2 deficiency by overexpression of WASL. Overexpression of wild type WASL in the TNK2 KO1 A549 cells led to increased EMCV virus infection (Figure 3C). Since constitutively active point mutants of WASL have been described (30, 31), (Figure 3-figure supplement1 B and C), we next tested whether overexpression of three such constructs in the TNK2 KO cells could further increase virus infection.. A higher level of complementation for all three mutants was observed compared to wild type WASL (Figure 3C). For NCK1, based on its reported binding to WASL and TNK2, we hypothesized that its function is to recruit WASL to TNK2, which could then activate WASL via phosphorylation (15, 32). Thus, we also tested whether constitutively active WASL could complement NCK1 KO. Constitutively active WASL fully rescued the NCK1 KO virus infection phenotype (Figure 3D and Figure 3-figure supplement1 D). Together, these data demonstrated that TNK2, WASL, and NCK1 are in a pathway to support picornavirus infection with WASL and NCK1 downstream of TNK2.

We further evaluated the interactions between TNK2, WASL and NCK1 using biochemical and biophysical assays. Previous studies had reported interaction between NCK1 and WASL through far Western and pull-down assay (12) and co-localization of TNK2 and NCK1 through protein overexpression (11). Fluorescent protein tagged forms of TNK2, WASL, and NCK1 were individually expressed in 293T cells. WASL and NCK1 both distributed homogenously throughout the cytoplasm, while TNK2 formed puncta in the cytoplasm (Figure 3-figure supplement2), which agreed with previous observations (21, 33). To examine potential co-localization, we co-expressed either mCerulean tagged TNK2 or mCerulean tagged WASL with mVenus tagged NCK1 in 293T cells, respectively. Since NCK1 is an adaptor protein, we reasoned that its interactions with binding partners would likely be relatively stable and therefore more readily detectable. Co-expression of mCerulean tagged TNK2 with mVenus tagged NCK1 re-localized NCK1 into the puncta observed by TNK2 expression alone (Figure 3-figure supplement2 and 3A). Co-expression of mCerulean tagged WASL with mVenus tagged NCK1 showed homogenous distribution in the cytoplasm and co-localization of these two proteins (Figure 3-figure supplement2 and 3A). Further analysis by quantitative FRET demonstrated that WASL and NCK1 had higher FRET efficiency in cells, while TNK2 and NCK1 also had significant, but lower, FRET efficiency (Figure 3-figure supplement 3A and 3B). Consistent with the FRET data, in co-transfected 293T cells, immunoprecipitation of FLAG tagged NCK1 pulled down HA tagged WASL more efficiently than it did Myc tagged TNK2 (Figure 3-figure supplement 3C). Altogether, these data demonstrate that TNK2 and WASL bind directly to NCK1.

### TNK2, WASL, and NCK1 affect a pre-replication step of the EMCV lifecycle

To begin dissecting the stage of the EMCV virus lifecycle impacted by TNK2, WASL, and NCK1, we transfected EMCV genomic RNA into the cells to bypass the early stages of the viral lifecycle. The TNK2 KO1, WASL KO, and NCK1 KO cells produced the same titers of EMCV virus as the control cells at 10 hours post transfection (Figure 4A) demonstrating that TNK2, WASL, and NCK1 are dispensable for EMCV replication and post-replication stages of the EMCV lifecycle. Thus, we concluded that all three genes act on an early stage of the viral lifecycle. As a second line of evidence, we determined the impact of adding the TNK2 kinase specific inhibitor Aim-100 before or at varying times after EMCV infection. While Aim-100 pre-treatment reduced EMCV infection, administration after virus inoculation had no effect (Figure 4B), consistent with a role for TNK2 at an early stage of virus infection.

**Figure 4.**
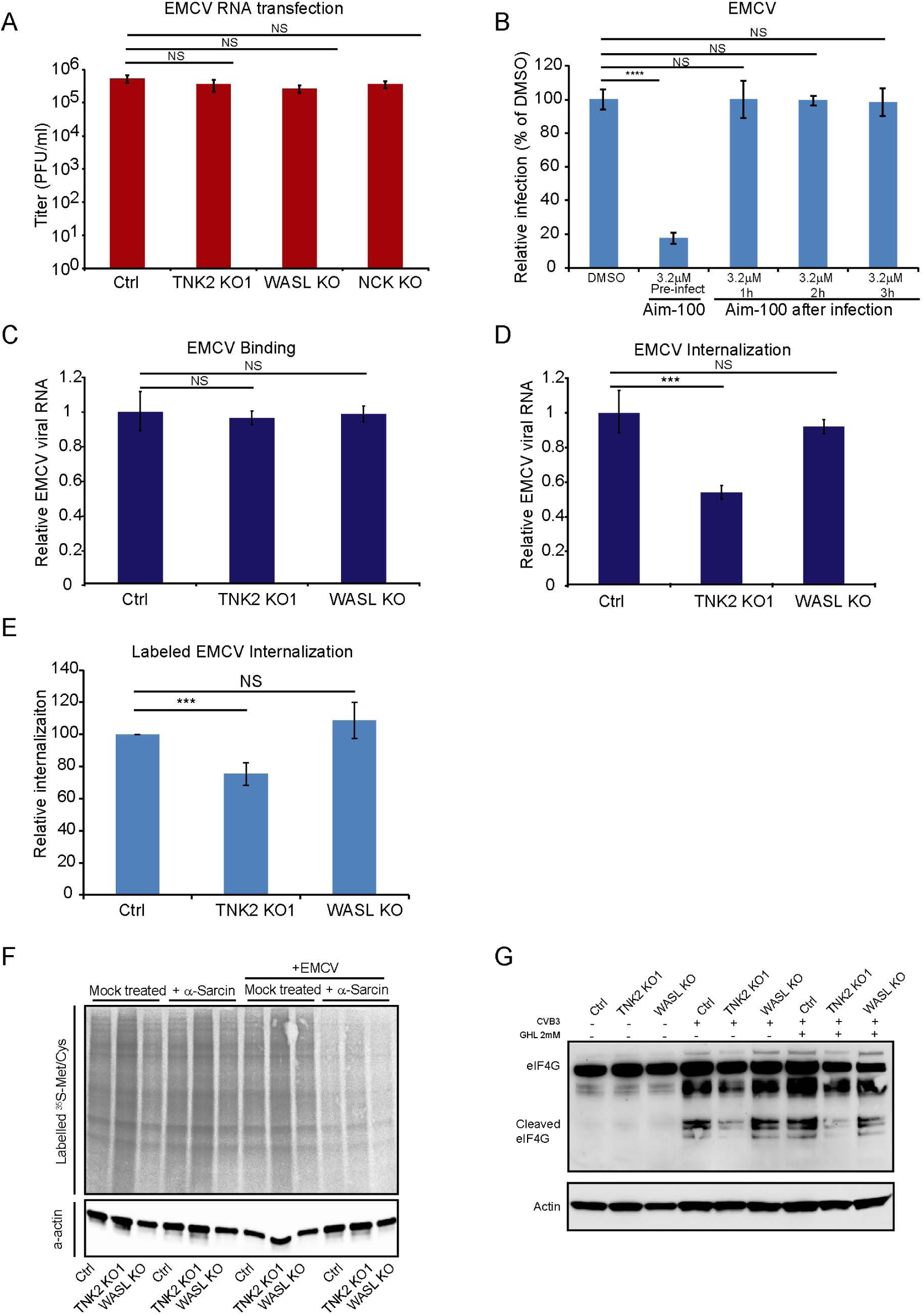
TNK2, WASL, and NCK1 function at an early stage of virus lifecycle. (A) EMCV released from viral RNA transfected Ctrl, TNK2 KO1, WALS KO, and NCK1 KO cells 10hours post transfection was quantified by plaque assay. (B) Time dependent addition of Aim-100 on EMCV infection on naïve A549 cells. A549 cells were treated with 3.2 µM Aim-100 at different time points before and after EMCV infection at an MOI of 1. EMCV positive cells were then quantified by FACS. (c) Quantification of EMCV virus binding on TNK2 KO1 and WASL KO cells by qRT-PGR expressed as relative change to Ctrl cell binding. (D) Quantification of EMCV virus internalization in TNK2 KO1 and WASL KO cells by qRT-PGR expressed as relative change to Ctrl cell internalization. (E) FACS quantification of labeled EMCV internalization in TNK2 KO1, WASL KO and Ctrl cells. (F) α-sarcin pore forming assay performed on TNK2 KO1, WASL KO and Ctrl cells. Translation was measured by phosphorimaging of 35S-methionine/ cysteine incorporation. (G) elF4G cleavage by GVB3 infection for 2 hours with or without 2mM guanidine hydrochloride. (A-E) Error bars represent standard deviation of three replicates. The data shown is representative of two independent experiments. ***: P<0.001, ****: P<0.0001, NS: not significant (P>0.05).

### TNK2, but not WASL functions in virus internalization

We examined the impact of TNK2 and WASL deletion on virus binding, virus internalization, pore formation, and viral RNA genome translocation. Following incubation at 4°C for one hour, no difference in virus binding was observed among control, TNK2 KO1, and WASL KO cells (Figure 4C), as assessed by qRT-PCR for EMCV genomic RNA. In contrast, after raising the temperature to 37°C for 30 minutes to allow for internalization of the bound virus particles and then trypsinization to remove uninternalized surface virus particles as described before (34, 35), there was a 46% reduction (P<0.001) of intracellular EMCV viral RNA in the TNK2 KO1 cells (Figure 4D). However, no difference in RNA levels was observed in WASL KO cells (Figure 4D). As a complementary approach, we evaluated internalization using fluorescently labeled EMCV virions (Figure 4-figure supplement A and B). 25% fewer EMCV positive TNK2 KO1 cells were observed compared to control cells (Figure 4E and Figure 4-figure supplement B), while no statistical difference was observed in the WASL KO cells (Figure 4E and Figure 4-figure supplement B).

We examined the ability of fluorescently labeled transferrin, a known cargo for clathrin mediated endocytosis, to be endocytosed in both TNK2 KO1 and WASL KO cells. Transferrin was internalized in TNK2 KO1 and WASL KO cells to the same level as in the control cells (Figure 4-figure supplement C and D), consistent with a previous report that knock down of TNK2 has no effect on transferrin endocytosis (36). To further check other general virus entry pathways, we also examined whether macropinocytosis was affected in these cells. FITC conjugated Dextran (7 KDa) is reported to be uptaken by macropinocytosis (24). No defect in uptake of Dextran was observed in TNK2 KO1 cells (Figure 4-figure supplement E and F). In contrast, the WASL KO cells had a 75% reduction in Dextran uptake (Figure 4-figure supplement E and F), which is consistent with the known dependency of macropinocytosis on actin polymerization (37). The Dextran macropinocytosis defect in the WASL KO appears to be unrelated to EMCV infection though since no reduction in either binding or internalization of EMCV was observed.

A step that is closely linked to internalization for picornaviruses is virus-induced pore formation, which is required for genome release into the cytoplasm. To assess pore formation, cells were treated with the membrane impermeable translation inhibitor, α-sarcin. Virus-induced pore formation will lead to translation inhibition and reduction of ^35^S methionine and ^35^S cysteine incorporation whereas in the absence of pore formation, translation will proceed at wild type levels (38). TNK2 KO1 and WASL KO cells showed no defect in virus pore formation during EMCV infection (Figure 4F). Although a clear defect in EMCV internalization exists in the TNK2 KO1 cells (Figure 4D and 4E), no defect in pore formation was observed. Since the internalization defect in TNK2 KO1 cells is not absolute, pore formation may still occur with sufficient frequency to inhibit translation. Alternatively, pore formation may not necessarily require prior internalization; for example binding of parainfluenza virus alone is sufficient to initiate pore formation at the cell membrane (39). For WASL, no defect in binding, internalization or pore formation was observed in the knockout cells, suggesting that the defect is downstream of these steps. Similarly, a recent report suggests that the human gene PLA2G16 acts at a step of the picornavirus lifecycle that is subsequent to pore formation (40).

To determine whether TNK2 or WASL plays any role in downstream viral RNA genome translocation after pore formation, we examined virus induced host protein cleavage after its entry. For some picornaviruses, immediately after translocation of its genome, translation of the incoming positive strand RNA leads to proteolytic cleavage of the host protein eIF4G to shut down host translation (41). Since this activity has not been described for EMCV we used CVB3, which is known to have this activity (42). eIF4G cleavage in presence or absence of a virus genome replication inhibitor guanidine hydrochloride showed no difference in the WASL KO cells as compared to the control cells, demonstrating that WASL does not impact this step of the viral lifecycle. In contrast, we observed a decrease of eIF4G cleavage in TNK2 KO1 cells, suggesting that TNK2 might play a role in this process. Alternatively, the reduced eIF4G cleavage level might simply reflect the reduced level of virus internalization in TNK2 KO1 cells (Figure 4G).

### Partial co-localization of TNK2 with labeled EMCV particles

TNK2 is present in endocytic vesicles and colocalizes with the early endosome marker EEA1 (21, 33, 36, 43). To visualize TNK2 subcellular localization, we expressed N-terminally GFP tagged TNK2 in the TNK2 KO1 cells. Two different patterns were observed in GFP positive cells: high TNK2 expression formed GFP aggregates inside the cytosol of cells in about 80% of the cell population, which has been described previously (21, 44), and low TNK2 expression inside the cytosol of cells in about 20% of the cell population(Figure 5-figure supplement A and B). Consistent with the native TNK2 rescue, N-terminally GFP tagged TNK2 expression in the TNK2 KO1 cells statistically increased virus infection from 17.2% to 22.0% of GFP transduced control cell infection (P<0.01, Figure 5-figure supplement D). The modest increase was comparable to that observed with ectopic expression of wild type TNK2. Next, we checked TNK2 localization in cells infected with fluorescently labeled EMCV. In TNK2 KO1 cells with high TNK2 expression that formed aggregates, no GFP-TNK2 co-localization with or in proximity to EMCV was observed. However, in cells with low TNK2 expression, labeled EMCV particles were observed in proximity to or surrounded by GFP-TNK2 (Figure 5A and Figure 5-figure supplement A), presumably in early endosomes. Further quantification demonstrated that about 40% of the EMCV particles were in proximity to GFP-TNK2 in those cells (Figure 5-figure supplement C).

**Figure 5.**
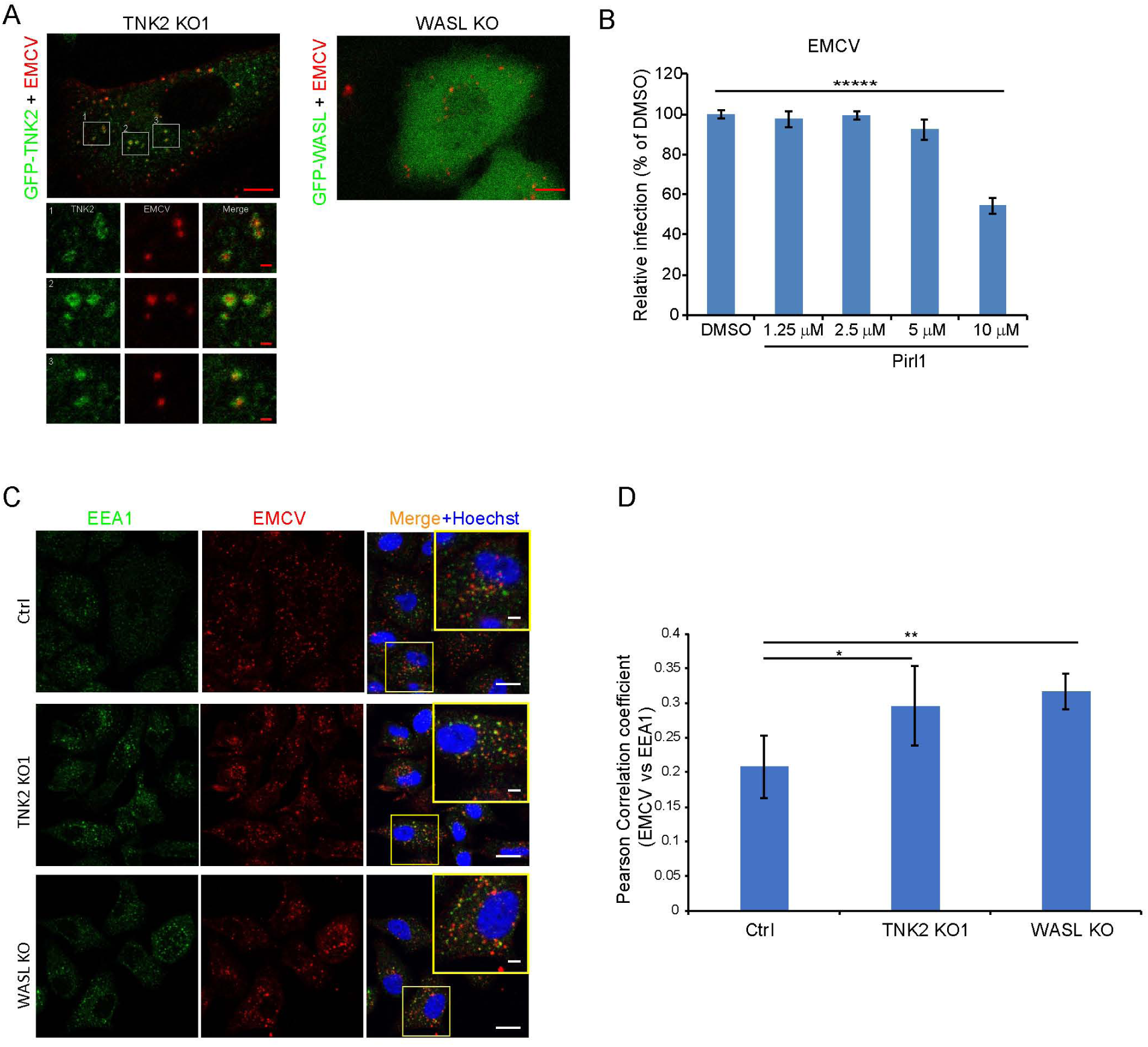
TNK2 mediates virus infection through endosomal trafficking pathways. (A) Confocal imaging of GFP-tagged TNK2 localization with fluorescently labeled EMCV virus in TNK2 KO1 cells and GFP-tagged WASL localization with fluorescently labeled EMCV virus in WASL KO cells. Scale bars represent 10µm. Individual channels of different insets were shown. Scale bars represent 2 µm. (B) FACS quantification of EMCV infection on pirl1 treated A549 cells at 10 hours post infection at an MOI of 1. (C) EEA1 staining of fluorescently labeled EMCV infected Ctrl, TNK2 KO1, and WASL KO cells. Scale bars represent 20 µm. Insets represent magnification of the boxed region. Scale bars represent 5 µm. (D) Quantification of Pearson correlation coefficient of EMCV and EEA1 colocalization in Ctrl, TNK2 KO1, and WASL KO cells infected with fluorescently labeled EMCV. (B, D) Error bars represent standard deviation of three replicates. The data shown is representative of two independent experiments. *: P<0.05, **: P<0.01.

WASL is recruited to vaccinia virus vesicles to promote its intracellular trafficking (23). To examine whether EMCV infection recruits WASL as observed for poxvirus, we expressed N-terminal GFP tagged WASL in WASL KO cells. The GFP tagged WASL was functional as it rescued EMCV infection (Figure 5-figure supplement E). Following infection with fluorescently labeled EMCV, no clear recruitment of GFP-WASL to fluorescently labeled EMCV was observed (Figure 5A).

### EMCV infection is CDC42 dependent but clathrin independent

Both TNK2 and WASL are activated by the upstream CDC42 kinase (11, 36, 45-47), which can be inhibited by pirl1 (24). Pirl1 treatment inhibited EMCV replication at a 10 µM concentration without cell toxicity (Figure 5B and data not shown), suggesting that CDC42 mediates EMCV infection.

Since a role for TNK2 in clathrin mediated endocytosis (CME) has been reported (21), we next examined whether the classic CME inhibitors such as dynasore (a canonical inhibitor of dynamin) or pitstop-2 (an inhibitor of clathrin) affect EMCV virus infection. Dynasore and pitstop-2 had no effect on EMCV infection while they inhibited VSV, a virus known to rely predominantly on CME for its entry, in a dose dependent fashion (Figure 5-figure supplement F and G). Together, these data indicate that EMCV infection in A549 cells is dependent on CDC42, which activates TNK2 and WASL, but does not require CME for its entry.

### EMCV virions accumulate in early endosomes in the absence of TNK2 or WASP

We next examined colocalization of EMCV with the early endosome marker EEA1. A time course analysis demonstrated that in control cells, the peak colocalization of EMCV with EEA1 occurred at 20 minutes post internalization (Figure 5-figure supplement H and I), which was followed by a decrease at 30 minutes post internalization. This result is consistent with a previous report that poliovirus entry peaks at 20 minutes post infection (4). In contrast, in both the TNK2 KO1 and WASL KO cells, higher levels of EMCV particles were retained in EEA1 containing vesicles at 30 minutes post internalization (Figure 5C and D, Figure 5-figure supplement I). Thus, both TNK2 KO1 and WASL KO cells are characterized by increased accumulation of EMCV particles in early endosomes.

### EMCV infection is dependent on the activation of WASL and its actin modulating function

The primary known effector function of WASL is nucleation of actin polymerization, which is mediated by binding of its C-terminal domain to the Arp2/3 complex and actin monomers (48). Different domains of WASL are critical for interaction with other proteins that can modulate WASL stability, activation, or function. Because ectopic expression of wild type WASL fully rescued EMCV infection in WASL KO cells, we tested the ability of a series of WASL domain deletion mutant constructs to rescue infection in WASL KO cells (Figure 6A). The basic region that binds with PIP2, the proline rich region that binds with SH3 domain containing proteins, the GTPase binding domain that interacts with CDC-42 kinase, and the acidic domain that is involved in actin binding and polymerization were all required for EMCV infection (Figure 6B and 6C). Interestingly, transduction of the N-terminal WH1 domain truncated mutant increased EMCV infection by 3-fold compared to wild type WASL transduction (Figure 6B). The WH1 domain is reported to bind with WASP interacting proteins (WIPs) to maintain WASL in an inactivated state (49); therefore, the WH1 domain truncation mutant should have increased actin polymerization activity. To further assess the WASL pathway’s role in EMCV infection, we treated cells with the Arp2/3 complex inhibitor CK-869. CK-869 treatment inhibited EMCV infection in a dose-dependent manner without cell toxicity (Figure 6D and data not shown). Altogether, these data demonstrated that WASL and its downstream actin pathway play important roles in EMCV infection.

**Figure 6.**
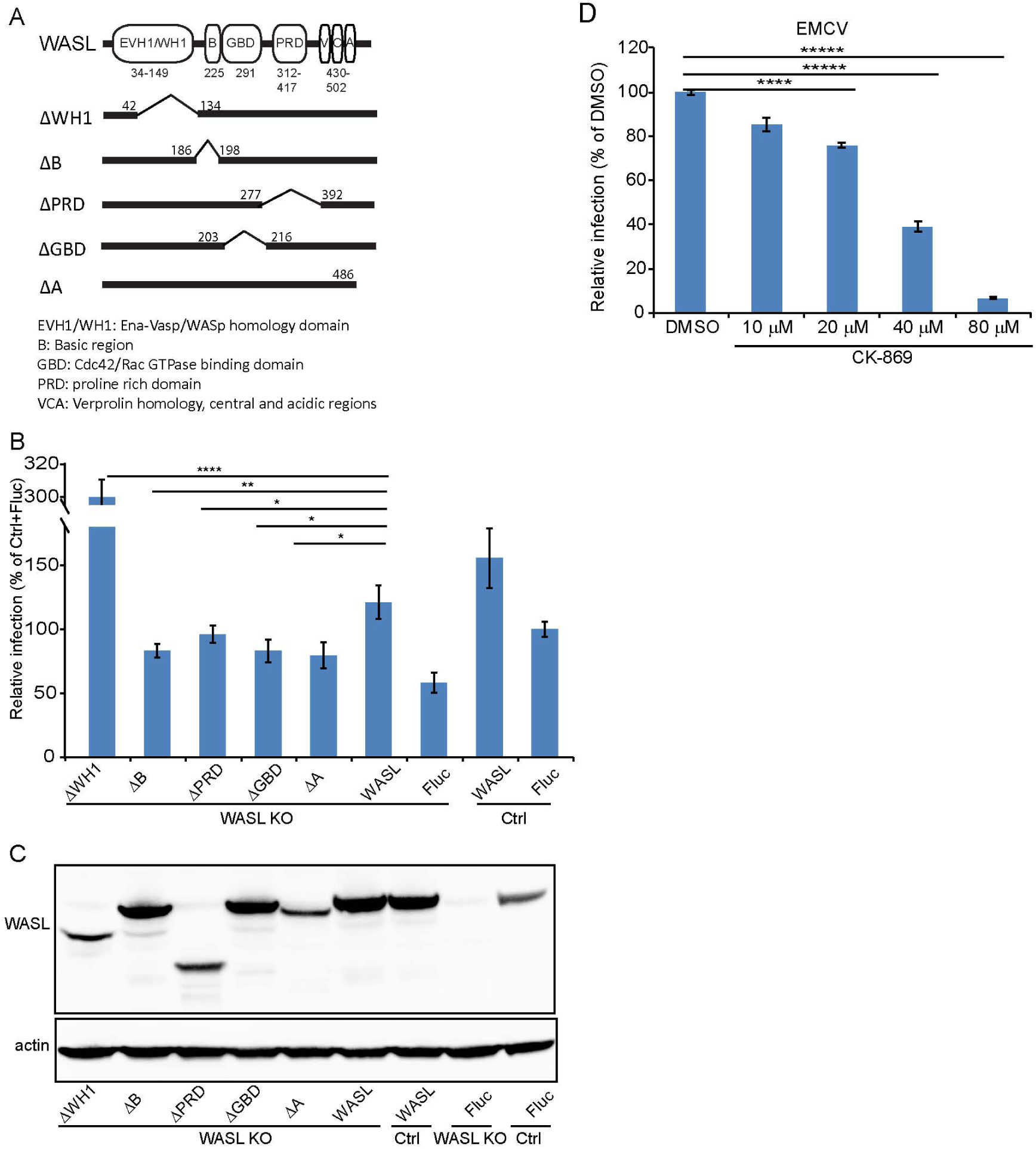
WASL activation and its actin modulation are critical for EMCV virus infection (A) Schematic representation of different WASL domain truncations. Each truncation is indicated by amino acid position on the constructs. (B) FACS quantification of EMCV infection in WASL KO cells transduced with different WASL domain truncations. (C) Western blot detection ofWASL domain truncation expression constructs in lentivirus transduced WASL KO cells. (D) CK-869 inhibition of EMCV infection on naïve A549 cells at 10 hours post infection at an MOI of 1. (B, D) Error bars represent standard deviation of three replicates. The data shown is representative of two independent experiments. *: P<0.05, **: P<0.01, ***: P<0.001, ****: P<0.0001.

### TNK2 is required for EMCV infection in an *in vivo* mouse model

To investigate TNK2’s role in EMCV infection *in vivo*, we generated a TNK2 knockout mouse that carried a 13Kb deletion of the murine TNK2 genomic locus, which eliminated all annotated TNK2 isoforms and splice variants (Figure 7A). In primary mouse lung fibroblast cells, complete ablation of TNK2 protein expression was observed by Western blot (Figure 7B). The number of cells infected by EMCV was reduced by 40% in the knockout cells (P<0.001, Figure 7C). *In vivo*, EMCV challenge with 10^7^ PFU by oral gavage resulted in greater survival of TNK2 knockout animals compared to wild type animals (P=0.0051, Figure 7D). Together, these data demonstrate a critical role for TNK2 in EMCV infection in an *in vivo* mouse infection model.

**Figure 7.**
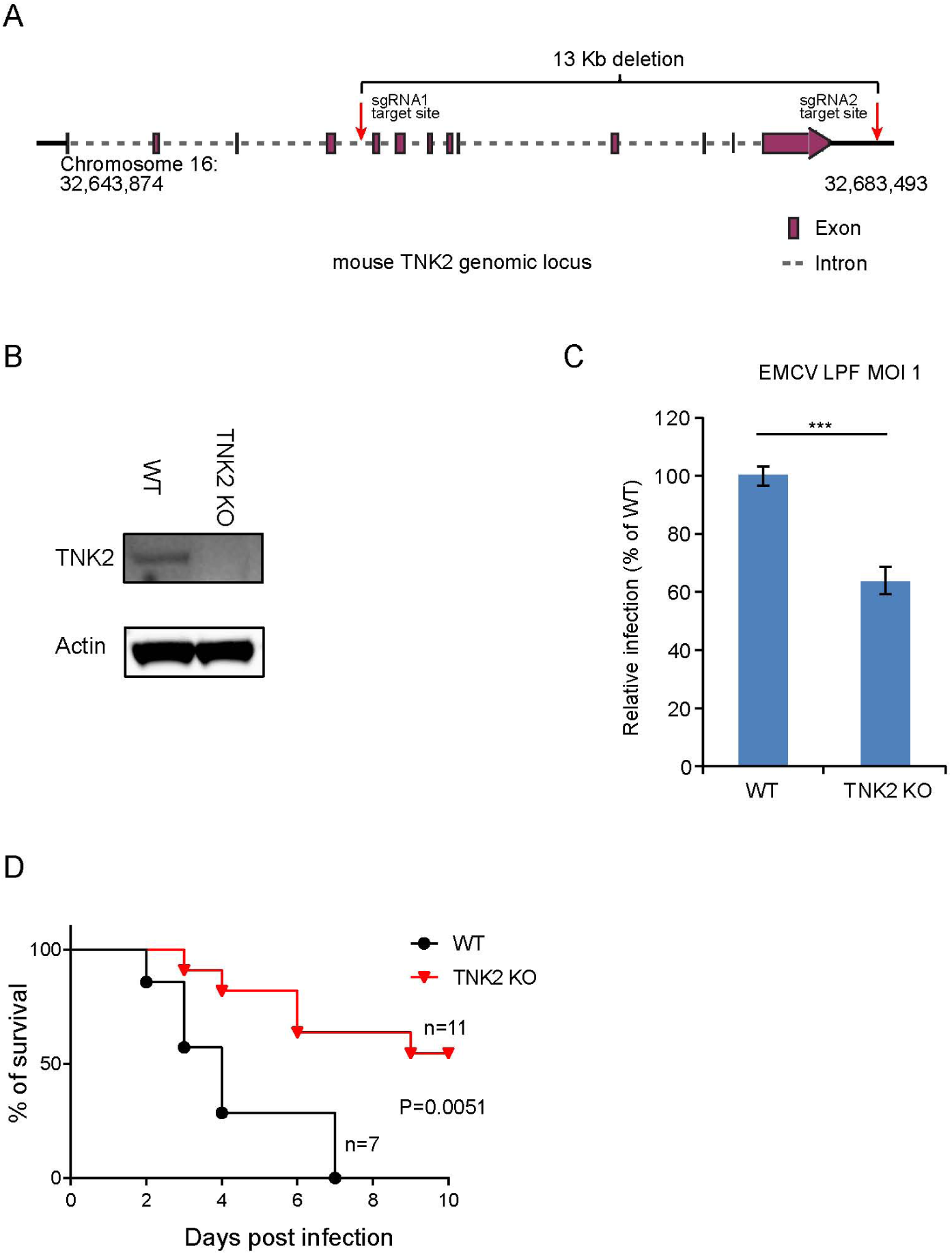
TNK2 is required for EMCV infection in vivo (A) Schematic representation of TNK2 knockout design by CRIPSR-Cas9 genome editing in mouse. Exon, intron and genomic position are indicated. (B) TNK2 expression in mouse primary lung fibroblast cells derived from TNK2 knockout and wild type animals. Cell lysates were analyzed by Western blot. (C) FACS quantification of EMCV infection in mouse primary lung fibroblast cells derived from TNK2 knockout and wild type animals 6 hours post infection at an MOI of 1. Error bars represent standard deviation of three replicates. The data shown is representative of two independent experiments. ***: P<0.001. (D) Survival curve of EMCV infection via oral gavage in TNK2 knockout and wild type mice. P=0.0051 by log-rank test.

## Discussion

We previously determined that the *C. elegans* genes *sid-3, viro-2*, and *nck-1* are essential for Orsay virus infection in *C. elegans* (10). In this study, we further asked whether their respective human orthologues TNK2, WASL, and NCK1 play any roles in mammalian virus infection. After screening a panel of mammalian viruses, we found that multiple picornaviruses, including EMCV, CVB3, enterovirus D68, and poliovirus, rely on TNK2, WASL, and NCK1 for infection in different cell lines. For TNK2, this represents the first evidence that it has any pro-viral function. Interestingly, poliovirus entry was previously shown to be dependent on an unknown kinase based on the use of a broad-spectrum tyrosine kinase inhibitor (4). One possibility is that TNK2 might be the kinase targeted in that study.

Our data represent the first demonstration that WASL and NCK1 are important for infection by picornaviruses such as EMCV, CVB3, enterovirus D68, and poliovirus. RNA transfection of EMCV into TNK2, WASL and NCK1 knockouts each yielded the same level of virus as the control demonstrating their role in an early stage of the EMCV lifecycle. For WASL, this is in contrast to its well-defined role in facilitating spread and transmission of vaccinia virus (22) suggesting that WASL is important for different lifecycle stages for different viruses. In addition to the known interactions between NCK1 and WASL, TNK2 has been reported to phosphorylate WASL (15). Consistent with these data, our genetic epistasis analysis and ectopic trans-complementation data (Figure 3A-D) demonstrated that these three genes function in a pathway in the context of picornavirus infection. Furthermore, biochemical and biophysical data demonstrated direct interaction of NCK1 with both TNK2 and WASL. Previous studies have established a clear role for CDC42 in regulation of TNK2 and WASL (47, 50), inhibition of CDC42 resulted in reduced EMCV infection.

We explored the mechanism by which TNK2 and WASL act regarding picornavirus infection. TNK2 KO1 cells were partially defective in EMCV internalization. In contrast, internalization was not affected in WASL KO cells, suggesting WASL functions in one or more steps downstream of virus internalization. In the absence of either TNK2 or WASL, EMCV particles continued to accumulate in early endosomes at 30 minutes post infection in contrast to control cells, demonstrating a role for both genes in endosomal trafficking. Notably, *sid-3*, the *C. elegans* orthologue of TNK2, is important for endosomal trafficking of RNA molecules (51). Thus, one possibility is that the delay in release from the early endosomes contributes to the reduction in EMCV infection in both TNK2 and WASL KO cells. However, it is uncertain whether this observed accumulation in early endosomes is directly related to the reduced levels of infection in these cells. In detailed studies of poliovirus, most of the RNA genome is released from the virus particles by 20 minutes post internalization (4). Thus, additional studies are necessary to determine whether the accumulated particles in the early endosomes still contain genomic RNA.

These data lead us to the following model (Figure 8). TNK2 is in a pathway upstream of WASL and NCK1. Because TNK2 deletion has a greater magnitude of impact than WASL or NCK1 deletion, TNK2 must act on one or more additional step of the viral lifecycle than WASL and NCK1. Consistent with this model, we demonstrated that TNK2 (but not WASL) is needed for virus internalization. In addition, we propose that TNK2 activates WASL (presumably, but not necessarily, by phosphorylation and in a fashion mediated by NCK1 binding), which then affects a subsequent early step of the EMCV lifecycle. Our data suggest that this step is linked to proper endocytic trafficking of the incoming viral particles, which accumulated in early endosomes in both TNK2 and WASL deficient cells to a greater extent than in control cells. The requirement for the WASL acidic domain, which is responsible for actin binding and nucleation, along with the sensitivity to the Arp2/3 inhibitor CK-869, demonstrates that this process is actin dependent. One possibility is that regulated actin polymerization may be necessary to deliver endocytic vesicles to specific subcellular locations necessary for productive infection.

**Figure 8.**
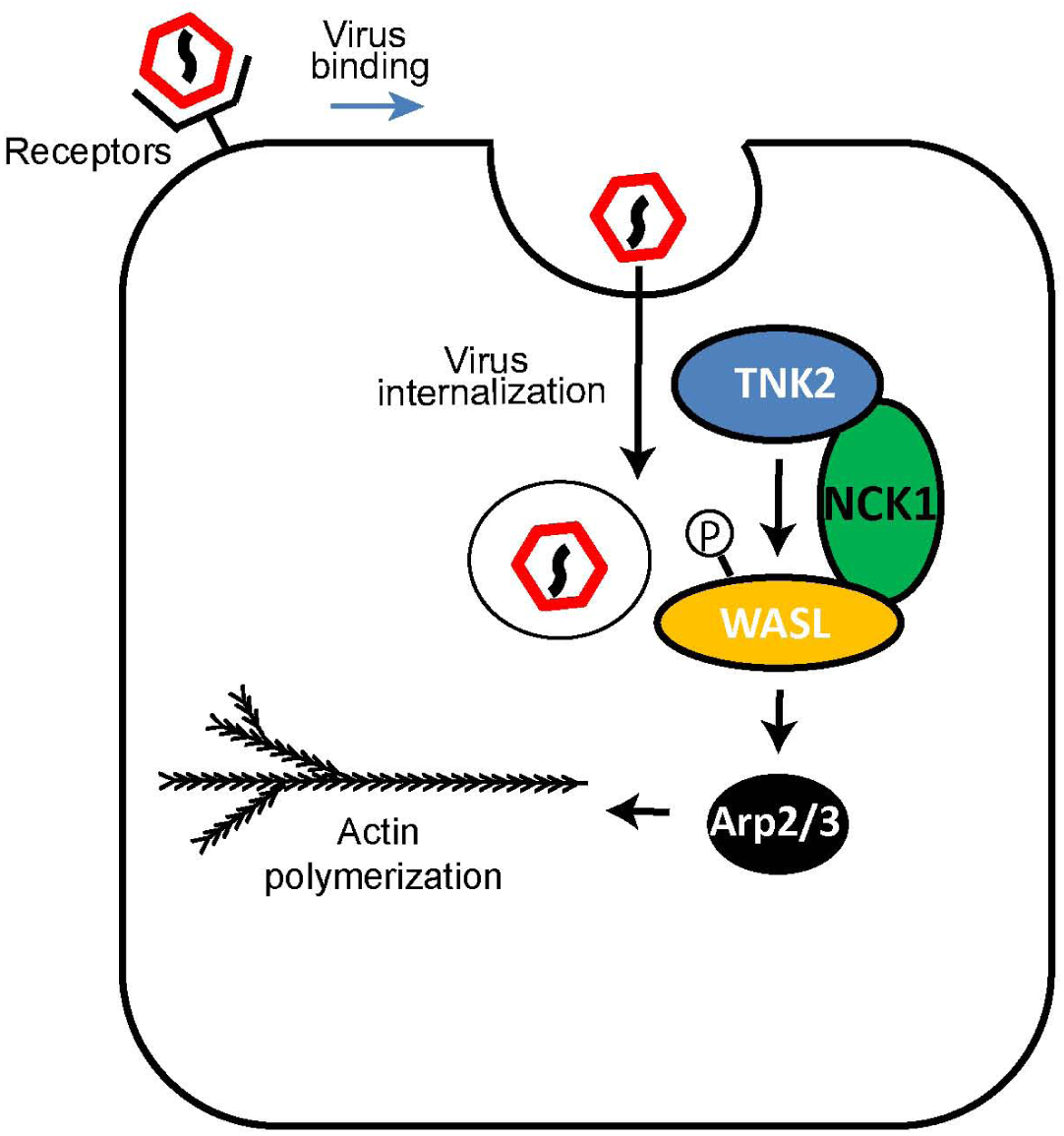
Model of TNK2, WASL and NCK1 function in picornavirus infection.

Actin is critical for many cellular processes and due to its essentiality is not a druggable target. In this study, we identified a TNK2 and actin-dependent pathway necessary to promote virus infection but does not appear to alter normal endocytosis. As cells lacking TNK2 and NCK1 are clearly viable, as are mice lacking TNK2 and NCK1 (52), these genes are potential antiviral targets that could be disrupted by small molecules without compromising overall actin biology and survival of the host.

It is becoming increasingly clear that there are additional, poorly defined stages of the picornavirus lifecycle. A recent genetic screen identified PLA2G16 as an essential gene required for an early stage of multiple picornaviruses at a step post binding, internalization, and pore formation (40). Our study similarly suggests that WASL also acts after these stages. Further study is needed to determine whether WASL acts at the same stage as PLA2G16 or at a distinct step.

All four picornaviruses we tested were dependent on TNK2 and WASL. In contrast, viruses in other families such as influenza A virus, parainfluenza 5 and adenovirus 5 were not affected by absence of these genes, suggesting that picornaviruses may specifically depend on these host factors. PIV5 is known to enter cells by direct fusion at the cellular membrane and is not believed to utilize endocytic vesicles (39). The observed lack of dependence of PIV5 on TNK2 or WASL is consistent with a potential role of TNK2 and WASL in modulating endocytic vesicle trafficking. In contrast, picornaviruses such as poliovirus, CVB3 and EMCV are generally thought to utilize clathrin independent endocytic vesicles for entry (5), while influenza A virus and adenovirus mostly rely on clathrin mediated endocytosis for their entry (2). The magnitude of the phenotypes did vary significantly among the tested picornaviruses, which is not surprising given the significant degree of divergence between these virus genomes and differences in their entry requirements. Nonetheless all four tested picornaviruses displayed different degree of dependence on this pathway. It will be important to test a wider range of picornaviruses to determine exactly which taxa within the *Picornaviridae* are most reliant on these genes. It will also be of great interest to test additional viruses from other families. The data suggesting a role of WASL in Lassa virus infection (24) supports the notion that at least some viruses in other virus families may also depend on one or more of these genes for infection.

In recent years, multiple studies of host factors critical for picornavirus infection have utilized haploid gene-trap screening or CRISPR screening in mammalian cell lines (40, 53, 54). These studies have identified multiple host genes required broadly for picornaviruses (40), as well as examples of genes required for specific picornaviruses. For instance, ADAM9 was discovered as a potential receptor for EMCV by CRISPR screening (54). Notably, the genes that we identified by starting with the *C. elegans* Orsay virus genetic screen, TNK2, WASL, and NCK1, were not identified as hits in any of the direct mammalian screens undertaken by others (40, 53, 54). Thus, these approaches provide complementary experimental strategies that, ideally in the long run, will lead to comprehensive understanding of host factors necessary for virus infection.

Our study of TNK2 was driven by our identification in *C. elegans* of *sid-3* as a host factor required for Orsay virus infection (10). *sid-3* was originally identified in a genetic screen for mutants defective in systemic spread of RNAi (51). That screening strategy also identified another *C. elegans* gene, *sid-1*, that is reported to transport dsRNA (55). A recent study demonstrated that the mammalian orthologue of *sid-1*, SIDT2, has an evolutionarily conserved function in transporting dsRNA that serves as a means of activating innate antiviral immunity in mice (56). Those studies provide an example of the value of using the *C. elegans* model system to guide insights into conserved functions with mammals. Here we demonstrated that TNK2, the human orthologue of *sid-3*, has an evolutionarily conserved function to facilitate virus infection in mammals. In *C. elegans*, Orsay virus infection is presumed to occur by fecal-oral transmission, as evidenced by its exclusive intestinal cell tropism (57). Deletion of *sid-3* reduces Orsay virus infection by > 5 logs *in vivo* in the *C. elegans* host (10). In the current study, infection of mice lacking TNK2 by oral gavage, the analogous route of infection as for Orsay virus infection of *C elegans*, led to increased survival of animals deficient in TNK2. These results clearly demonstrate the functional parallels between *sid-3* and TNK2 in their respective hosts *in vivo*. Thus, our study has identified another example wherein novel insight in mammalian biology emerge from initial discoveries in model organism studies.

## Material and methods

### Cell culture and viruses

A549 cells were cultured and maintained in DMEM supplemented with 25 mM HEPES, 2 mM L-glutamine, 1X non-essential amino acids, 10% Fetal bovine serum (FBS) and 100 u/ml antibiotics (penicillin and streptomycin). 293T (human embryonic kidney), BHK-21 (Baby hamster kidney), RD (rhabdomyosarcoma) and Hela cells were cultured and maintained in DMEM with 10% FBS. Haploid cells (HAP1) were cultured and maintained in IMDM with 10% FBS. Mouse primary lung fibroblasts were isolated by lung digestion as described (58). Primary lung fibroblasts were derived from a TNK2 homozygous knockout mouse and a homozygous wild type littermate and were cultured in DMEM with 20% FBS. Viruses were obtained from the following: EMCV VR-129 strain (Michael Diamond), coxsackie virus B3 Nancy strain (Julie Pfeiffer), poliovirus Mahoney strain (Nihal Altan-Bonnet), influenza A virus WSN strain (H1N1) (Adrianus Boon), adenovirus A5 (David Curiel), enterovirus D68 (ATCC), parainfluenza virus (Robert A. Lamb), GFP-EMCV and GFP-CVB3 (Frank J. M. van Kuppeveld). EMCV was amplified on BHK-21 cells; CVB3 and polio viruses were amplified on HeLa cells; and enterovirus D68 was amplified on RD cells.

### Reagents and antibodies

α-sarcin was purchased from Santa Cruz. ^35^S methionine and ^35^S cysteine were purchased from Perkin Elmer. Alexa Fluor™A647 succinimidyl ester was purchased from ThermoFisher Scientific. Inhibitors were purchased from commercial vendors as follow: Aim-100 (Apexbio), Wiskostatin (Sigma), Dynasore (Sigma), Pitstop-2 (Sigma), CK-869 (Sigma), Pirl1 (Hit2leads). Anti-EMCV mouse polyclonal antibodies were provided by Michael Diamond. Anti-poliovirus antibodies were provided by Nihal Altan-Bonnet. Other antibodies were obtained from commercial vendors as follows: Anti-coxsackie virus B3 antibodies (ThermoFisher), Anti-adenovirus A5 antibodies (ThermoFisher), Anti-influenza A virus NP antibodies (Millipore), Anti-TNK2 (A12) (Santa Cruz), Anti-WASL (Abcam and Sigma), Anti-actin, clone C4 (Sigma), Anti-NCK1 (Millipore), enterovirus D68 Ab (GeneTex), Anti-HA (ThermoFisher), Anti-Flag (GenScript), Anti-c-Myc (Invitrogen), Anti-double stranded J2 antibodies (Scicons).

### Plasmid constructions

Single guide RNA (sgRNA) oligonucleotides were synthesized by Integrated DNA Technologies (IDT). sgRNA oligoes were annealed and cloned into Lenti-CRISPR V2 plasmid digested by BsmBI. TNK2 (pReceiver-TNK2) and WASL (pReceiver-WASL) ORF clones were obtained from GeneCopoeia. NCK1 (pcDNA-NCK1) ORF clone was obtained from GenScript. Mutations in the sgRNA binding sites were introduced by site directed mutagenesis (Stratagene) according to the manufacturer’s protocol. TNK2, WASL and NCK1 were subcloned into a Lentivirus expression vector pFCIV digested with Asc1 and Age1. The FRET control plasmid C5V was obtained from Addgene. mVenus cassette was subcloned into pcDNA-NCK1 and pReceiver-WASL. mCerulean cassette was subcloned into pReceiver-TNK2. Myc tagged TNK2 and HA tagged WASL was generated by annealing of oligonucleotides and then subcloned into the expression vector. GFP tagged TNK2 and WASL were subcloned into a tetracycline promoter driven Lentivirus vector pCW57. Constitutive active WASL constructs were generated by site-directed mutagenesis. Domain truncation were generated by overlapping PCR. In brief, to generate pFCIV-WASL ΔWH1, pFCIV-WASL ΔB, pFCIV-WASL ΔPRD, pFCIV-WASL ΔGBD, and pFCIV-WASL ΔA, WASL fragments were amplified from the start of the gene to the start of the truncation and from the end of the truncation to the end of the gene, and the fragments were then joined by overlapping PCR. The resulting product was digested by AgeI and AscI and ligated into Lentivirus expression vector pFCIV that were cut by the same restriction enzymes.

### Lentivirus production and cell transduction

800 ng of Lenti CRISPR V2 plasmids or pFCIV plasmids or pCW57 plasmids were transfected with 800 ng pSPAX2 and 400 ng pMD2.G into 293T cells using Lipofectamine 2000 according to the manufacturer’s protocol. Cell culture supernatant was harvested two days post transfection and stored at −80°C. For transduction, A549 cells were seeded one day before transduction. Cells were spin transfected with lentivirus and 8 µg/ml of polybrene at MOI 5. Two days after transduction, cells were passaged and either selected under corresponding antibiotics or fluorescently sorted through flow cytometry.

### CRISPR genome editing

A549 naïve cells were transduced by lentiviruses that express the corresponding sgRNA and Cas9 protein. Transduced cells were passaged two days later and then selected with 2 µg/ml puromycin for 7 days. Cells were passaged once during this selection. Clonal selection was performed through limiting dilution in 96-well plates. After two weeks, single cell clones were picked and expanded in 24-well plates. The desired genome editing was identified by a restriction enzyme digestion-based genome typing assay. Genome edited clonal cells were further sequenced by Sanger sequencing to define the precise genome editing event. Detection of a homozygous 28bp (base pair) deletion allele of TNK2 defined TNK2 KO1 and detection of a 25bp deletion allele and 137bp deletion allele of TNK2 defined TNK2 KO2. Identification of a 1bp insertion allele and a 2 bp deletion allele of WASL defined WASL KO. An insertion allele of 94 bp in NCK1 was observed and this defined NCK1 KO. TNK2 KO1, WASL KO and NCK1 KO were used in all experiments except places specified using other cells.

For homologous template directed DNA repair through CRISPR genome editing, assembled CRISPR RNP were transfected into A549 cells with single stranded oligodeoxynucleotide (ssODN) as described (59). In brief, Alt-R® S.p. Cas9 nuclease 3NLS protein, Alt-R® CRISPR-Cas9 crRNA designed to target the deleted TNK2 KO1 genomic region, Alt-R® CRISPR-Cas9 ATTO™ 550 tagged tracrRNA and ssODN were purchased from IDT. A549 cells were seeded into 12-well plates one day before transfection. Equal molar crRNA and tracrRNA were annealed by heating to 95 °C for 5 minutes and then cooled to room temperature. RNP was assembled by combing equal molar ratio of annealed cr-tracrRNA with Cas9 nuclease protein in opti-MEM. RNP was then transfected with ssODN into A549 cells by lipofectamine CRISPR-MAX according to the manufacturer’s protocol. 24 hours after transfection, ATTO™ 550 positive cells were FACS sorted individually into 96-well plates. Single cell colonies were expanded one week after sorting and genotyped with a restriction enzyme based genotyping assay. Template directed DNA repair was finally confirmed by both Sanger and PCR product deep sequencing.

For generation of gene double or triple knockout cells, the same CRISPR RNP transfection method was used as homologous template directed DNA repair using corresponding crRNA designed to target genes with ssODN omitted. A double knockout of NCK1 and WASL was generated by CRISPR-Cas9 genome editing of the NCK1 locus in the WASL KO cells. A double knockout of WASL and TNK2, a double knockout of NCK1 and TNK2, and a triple knockout of WASL, NCK1 and TNK2 were generated by CRISPR-Cas9 genome editing of the WASL locus and NCK1 locus in the TNK2 KO1 cells. Genome editing events were screened by a restriction enzyme digestion-based genotyping assay.

### EMCV virus labeling and infection for imaging

EMCV was amplified in BHK-21 cells and purified according to previous publications (4, 40). Virus labeling was performed with Alexa Fluor™A647 succinimidyl ester in a 1:10 molar ratio and was then purified through Nap-5 desalting column (GE Healthcare). Labeled viruses were aliquoted and stored at −80°C. For EMCV entry imaging analysis, A549 cells were seeded at 3000 cells per well in an 18-well IBD imaging slide chamber one day before infection. The next day, cells were washed once with serum free DMEM and then inoculated with labeled EMCV virus at an MOI of 20 on ice for one hour. Cells were washed three times with ice-cold PBS after on ice binding. Cells were then fixed with 4% paraformaldehyde for 15 minutes at room temperature or cells were switched to 37°C incubation with complete medium for internalization. After incubation with complete medium for 30 minutes, cells were then washed once with PBS and fixed with 4% paraformaldehyde. Fixed cells were mounted in IBD mounting medium for image analysis. For GFP-TNK2 and GFP-WASL localization with labeled EMCV virus imaging, live cell experiment was performed in an 18-well IBD imaging slide in a temperature-controlled imaging chamber.

### EMCV virus binding and internalization assay

A549 cells were seeded at 1×10^5^ cell per well in a 24-well plate one day before the assay. Cells were chilled on ice for 30 minutes and then washed with ice-cold DMEM before inoculation. EMCV was diluted in ice-cold DMEM with 0.1% BSA and then inoculated at an MOI of 20 in 250 µl of DMEM per well of 24-well plate. Viruses were allowed to bind on ice for one hour. For virus binding experiments, cells were washed three times with ice-cold PBS and then lysed in 350 µl Trizol reagent. For virus internalization assay using trypsinization, experiments were performed as described previously for WNV and AAV (34, 35). In brief, after one hour of binding, the virus inoculum was removed and pre-warmed complete medium was added onto cells. Cells were incubated in a 37°C water bath for 30 minutes to allow for virus internalization. After incubation, cells were washed three times with ice-cold PBS and then trypsinized for 6 minutes to remove surface bound virus. Trypsinized cells were then washed again three times and then spin at 300g for 5 minutes. Cell pellets were lysed in 350 µl Trizol reagents. RNA was extracted using the 96-well Zymo RNA easy column extraction according to the manufacturer’s protocol. Viral RNA was quantified by one step reverse transcription quantitative real-time PCR with an EMCV assay probe (Primers: forward 5’-CGATCACTATGCTTGCCGTT-3’; reverse 5’-CCCTACCTCACGGAATGGG-3’; Taqman probe 5’ FAM-AGAGCCGATCATATTCCTGCTTGCCA-3’). Fold change was converted from delta delta Ct of an internal control assay by RPLP0 (Ribosomal protein lateral stalk subunit P0). For labeled EMCV, internalization was performed the same way, after trypsinization and washing, cells were fixed in 4% paraformaldehyde for 15 minutes at room temperature and then washed twice with P2F (PBS with 2% Fetal bovine serum). Cells were finally resuspended in P2F and analyzed by MACS flow cytometry.

### FACS assay

A549 cells were seeded one day before infection into 96-well plates. Approximately 16 hours after seeding, cells were infected by EMCV at an MOI of 1. One hour after infection, the inoculum was removed and cells were cultured in DMEM with 2% FBS. 10 hours post infection, cells were trypsinized and fixed with 4% paraformaldehyde. Fixed cells were then permeabilized with perm buffer (1g Saponin, 10ml HEPES, 0.025% Sodium Azide in 1L HBSS) for 15 minutes. After permeabilization, cells were incubated with primary antibodies for one hour and then washed twice before incubation with fluorescently conjugated secondary antibodies. After one hour of secondary antibody incubation, cells were washed three times with perm buffer and then resuspended with 70 µl of FACS buffer P2F (PBS with 2% fetal bovine serum). Infected cells were then analyzed and quantified through MACS flow cytometry (Miltenyi Biotec). FACS analysis of infection by CVB3, polio, enterovirus D68, adenovirus and influenza virus on either A549 or Hap1 cells were performed the same as EMCV infection, except that cells were harvested 8 hours post infection for CVB3, adenovirus, enterovirus D68 and influenza virus, and 6 hours post infection for poliovirus.

### Multistep growth analysis for EMCV and polio virus

For multistep growth analysis, A549 or Hap1 cells were infected by EMCV at an MOI of 0.01. One hour after inoculation, cells were washed 5 times with serum free DMEM and were then cultured in DMEM or IMDM with 2% FBS. Culture supernatant was collected at time 0, 6, 12, 24, 36 and 48 hours post infection. Viruses released in the culture supernatant were titrated on BHK-21 cells by plaque assay. For polio virus multi-step growth titration, A549 or Hap1 cells were infected by virus at MOI 0.01 and culture supernatant were collected. Released viruses were titrated on HeLa cells by plaque assay.

### EMCV genomic RNA transfection

EMCV genomic RNA was extracted from pelleted virions by phenol chloroform extraction. For RNA transfection, A549 cells (gene knockouts and control) were seeded into 24-well plate at 1×10^5^ cells per well. 16 hours after seeding, cells were transfected with 1.6 µg EMCV RNA by Lipofectamine 3000 according to the manufacturer’s protocol. Cell culture supernatant was collected at 10 hours post transfection and was then titrated on BHK-21 cells by plaque assay.

### Co-immunoprecipitation and Western blot

Flag C-terminal tagged NCK1 was transfected with HA N-terminal tagged WASL or with Myc N-terminal tagged TNK2 into 293T cells. 48 hours after transfection, cells were lysed in IpLysis buffer (Invitrogen). Protein concentrations were quantified by BCA assay. 500 µg protein lysates were incubated with 2 µg of anti-flag antibodies at 4°C for overnight for immunoprecipitation. After antibody binding, the immuno-complex was incubated with protein A/G magnetic beads for one hour at room temperature. The beads were washed and proteins bound to antibodies were eluted according to the manufacture’s protocol (Invitrogen). Protein samples were prepared with 4X NuPAGE sample buffer (Invitrogen) and then resolved on 4-12% NuPAGE gel. Proteins were transferred to a PVDF membrane and then blocked with 5% skim milk. Primary antibodies were incubated in 5% milk for overnight at 4°C. After 3 washes with PBST (PBS with 0.3% of tween-20), corresponding HRP conjugated secondary antibodies were incubated in 5% milk at room temperature for 1 hour. The membrane was washed 5 times with PBST and was then developed with chemiluminescent substrate for 5 minutes and then imaged using a Bio-Rad chemiluminescence imager.

### α-Sarcin pore forming assay

A549 cells were seeded at 3.6 × 10^4^ per well of a 48-well plate. The next day, cells were washed once with PBS and then incubated with methionine and cysteine free DMEM for one hour. Subsequently, cells were inoculated with EMCV virus at an MOI of 50 on ice for one hour. Inoculum were then removed and cells were washed three times with ice cold PBS. Cells were then incubated with 100 µg/ml α-sarcin diluted in methionine and cysteine free DMEM for 90 minutes at 37°C. After α-sarcin treatment, cells were pulsed with 7.4 MBq ^35^S methionine and ^35^S cysteine in DMEM for 20 minutes. Next, cells were washed four times with ice cold PBS and then lysed with SDS sample buffer. Proteins were resolved on 7.5% SDS-PAGE gel and then dried on a Bio-Rad gel dryer. Dried gels were exposed to a phosphorimager overnight and then scanned by Fuji IFA imaging system. As an internal loading control, a parallel SDS-PAGE gel was ran and then proteins were transferred to PVDF membrane and blotted with anti-actin antibodies.

### Immunofluorescence assay

For EMCV co-localization with EEA1 marker, A549 cells were seeded at 3000 cells per well in an 18-well IBD imaging slide chamber one day before infection. The next day, cells were washed once with serum free DMEM and then inoculated with labeled EMCV virus at an MOI of 20 on ice for one hour. Cells were washed three times with PBS after on ice binding. Cells were then switched to 37°C incubation with complete medium for internalization at indicated time. After incubation with complete medium for different time, cells were then washed once with PBS and fixed with 4% paraformaldehyde for 10 minutes. Fixed cells were then permeabilized with 0.2% saponin for 10 minutes at room temperature and subsequently blocked with IFA blocking buffer (10% goat serum, 0.05% saponin in PBS) at room temperature for 30 minutes. Cells were then incubated with EEA1 antibodies at 1:100 dilution in blocking buffer overnight with shaking. Cells were washed three times with PBS and then incubated with secondary antibodies at room temperature for 1 hour. After incubation, cells were washed once with PBS and then incubated with Hoechst at 1:1000 dilution for 10 minutes at room temperature. Cells were washed three time with PBS and mounted in IBD mounting medium for image analysis. For dsRNA immunostaining in EMCV infected cells, the same procedure was performed as EEA1 immunostaining except that 0.1% Triton was used instead of Saponin.

### Confocal imaging and FRET analysis

Cells on slides or in an IBD imaging slide chamber were examined on a Zeiss airy scan confocal microscope (LSM 880 II). A Plan Apochromat 63X, 1.4-numerical-aperture oil objective lens (Carl Zeiss, Germany) was used to image labeled virus infection. For FRET analysis, 293T cells were seeded on coverslips and transfected with FRET pairs, mCerluean tagged TNK2 with mVenus tagged NCK1 and mCerulean tagged WASL with mVenus tagged NCK1 respectively. 24 hours post transfection, cells were fixed in 4% paraformaldehyde and mounted on slides. Cells were imaged on a Zeiss airy scan confocal with a Plan Apochromat 100X, 1.4-numerical-aperture oil objective lens. Acceptor photo bleach was performed with 80% laser intensity of the imaging channel. Images were taken before and after photo bleach and FRET efficiency were calculated after image acquisition on Zen pro software (Carl Zeiss, Germany). For quantification of EMCV colocalization with EEA1, image analysis was performed using Volocity software V6.3 (PerkinElmer).

### Transmission electron microscopy

EMCV was incubated with A549 cells (wild type, TNK2 KO and WASL KO) at an MOI of 20 for 1 hour on ice. Cells were then washed with serum free DMEM and incubated with A549 culture media (2% FBS) for 6 hours. Infected cells were washed with PBS and fixed with 2% paraformaldehyde, 2.5% glutaraldehyde (Polysciences Inc., Warrington, PA) in 100 mM cacodylate buffer for 1 hour at room temperature. Next, cells were scraped from plates using a rubber cell scraper and cell pellets were embedded in agarose. Agarose embedded cell pellets were post-fixed in 1% osmium tetroxide (Polysciences Inc.) for 1 hour, then rinsed extensively in dH20 prior to *en bloc* staining with 1% aqueous uranyl acetate (Ted Pella Inc., Redding, CA) for 1 hour. Following several rinses in dH20, samples were dehydrated in a graded series of ethanol and embedded in Eponate 12 resin (Ted Pella Inc.). Sections of 95 nm were cut with a Leica Ultracut UCT ultramicrotome (Leica Microsystems Inc., Bannockburn, IL), stained with uranyl acetate and lead citrate, and viewed on a JEOL 1200EX transmission electron microscope (JEOL USA, Peabody, MA) equipped with an AMT 8 mega-pixel digital camera (Advanced Microscopy Techniques, Woburn, MA).

### *In vivo* infection experiments

Animal experiments were conducted under the supervision of Department of Comparative Medicine at Washington University in St. Louis. All animal protocols were approved by the Washington University Institutional Animal Care and Use Committee (Protocol #20170194 and #20180289). TNK2 knockout mice were generated in the C57BL/6 background by CRISPR-Cas9 genome editing at the Genome Engineering and iPSC Center (GEiC) at Washington University. All animals were housed in the pathogen free barrier. Age-matched animals with mixed gender (6-8 weeks old, 5 females and 6 males for TNK2 knockout, 2 females and 5 males for wild type littermates) were infected with 1×10^7^ PFU of EMCV (two doses at day 0 and day 1) via oral gavage according to previous publication (60). Infected mice were monitored for at least 10 days for all experiments.

### Statistical analysis

For statistical analysis, Student T-test was performed on the average values from three replicates. All data shown with statistics are representative of at least two independent experiments. Log-rank test was done in Graphpad Prism V7. Statistical significance are indicated as below: NS: no statistical significance, *: P<0.05, **: P<0.01, ***: P<0.001, ****: P<0.0001, *****: P<0.00001.

## Acknowledgement

This work was supported in part by National Institutes of Health R01 AI134967.

We thank Adrianus Boon, Skip Virgin, Celeste Morley and John Cooper for helpful discussions. We thank Rong Zhang and Michael S. Diamond for access to FACS instrumentation. FRET experiments were performed at the Washington University Center for Cellular Imaging (WUCCI) supported by Washington University School of Medicine, The Children’s Discovery Institute of Washington University and St. Louis Children’s Hospital (CDI-CORE-2015-505) and the Foundation for Barnes-Jewish Hospital (3770). We thank Wandy Beatty for assistance with electron microscopy and laser scanning confocal. We thank the Genome Engineering and iPSC Center (GEiC) at the Washington University in St. Louis for their sgRNA validation and genotyping services for generating the TNK2 knockout mouse. We thank the flow cytometry core at Department of Pathology and Immunology, Washington University School of Medicine for assisting cell sorting. We thank Tim Schaff and Darren Kreamalmeyer for assistance with mouse breeding.

## Author Contributions

H.J. performed the *in vitro* and *in vivo* experiments, including the primary CRISPR/Cas9 knockout of TNK2, WASL, validation in different cells with different viruses, gene rescue, RNA genome transfection, virus binding, internalization, pore-formation, host factor cleavage, FRET, co-immunoprecipitation, co-localization assays, *etc*. C.L. generated expression clones, part of the double knockout clones, performed the constitutive WASL rescue, domain analysis of WASL, and generated figure 3A-D and figure 6A-C. S.T. generated NCK1 KO cells, conducted the experiments in figure 2F-H, and performed quantitative image analysis. H. J. and D. W. designed the experiments and performed data analysis. H.J. wrote the initial draft of the manuscript, with D. W. editing and the other authors contributing to the final paper.

---

**Figure 1-figure supplement1.**
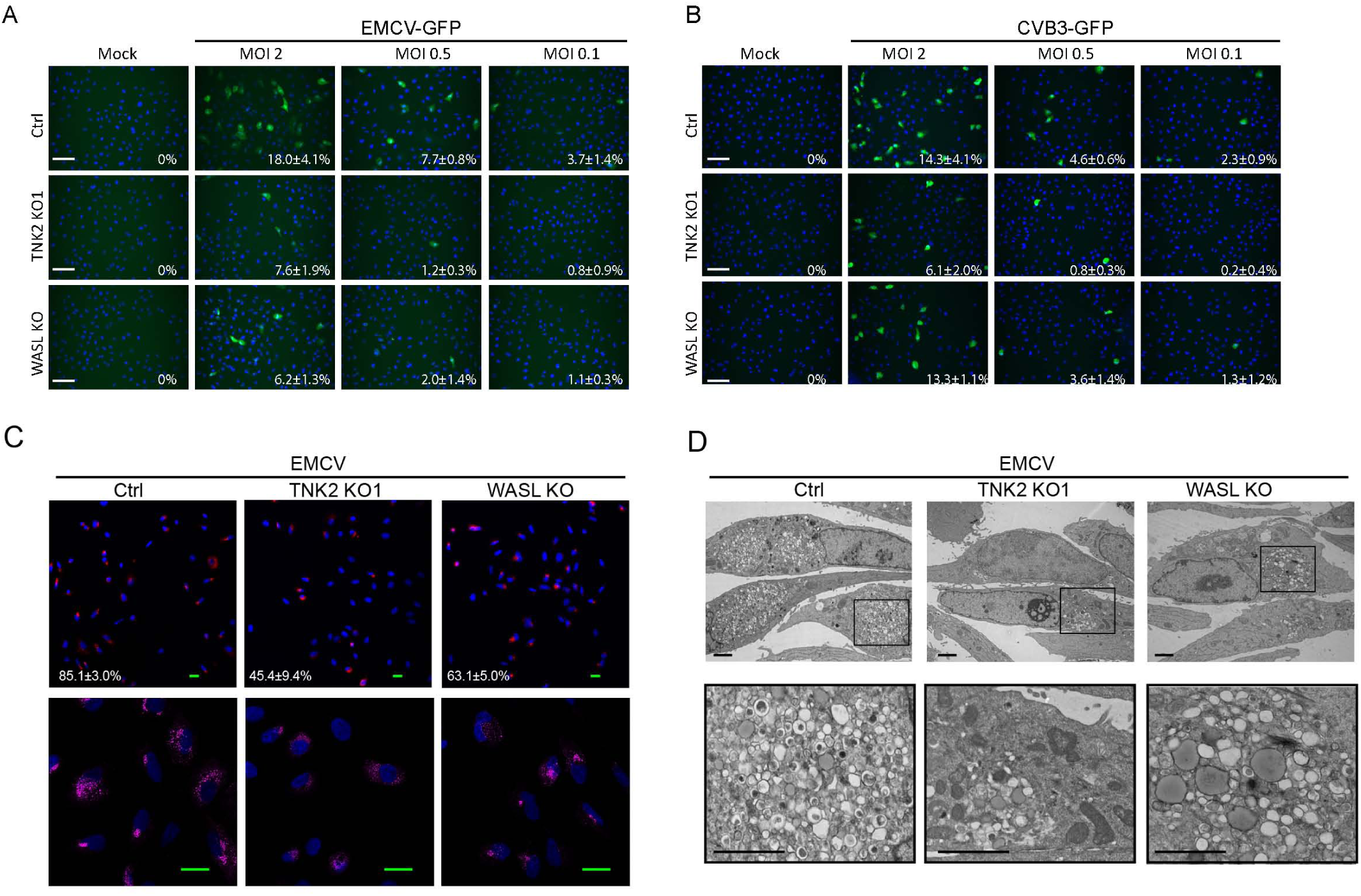
TNK2 and WASL are critical for multiple picornavirus infection on A549 cells. (A) EMCV-GFP infection on Ctrl, TNK2 KO1 and WASL KO A549 cells at MOI 2, 0.5 and 0.1. Percentage of positive cells were quantified (values denote mean±s.d., n= 3). Scale bars represent 20 µm. (B) CVB3-GFP infection on Ctrl, TNK2 KO1 and WASL KO A549 cells at MOI 2, 0.5 and 0.1. Percentage of positive cells were quantified (values denote mean±s.d., n= 3). Scale bars represent 20 µm. (C) dsRNA immunostaining of EMCV infection on Ctrl, TNK2 KO1 and WASL KO A549 cells. Percentage of positive cells were quantified (values denote mean±s.d., n= 3). Scale bars represent 10 µm. (D) Electron microscopy detection of EMCV replication complex on Ctrl, TNK2 KO1, and WASL KO A549 cells. Scale bars represent 2 µm.

**Figure 1-figure supplement2.**
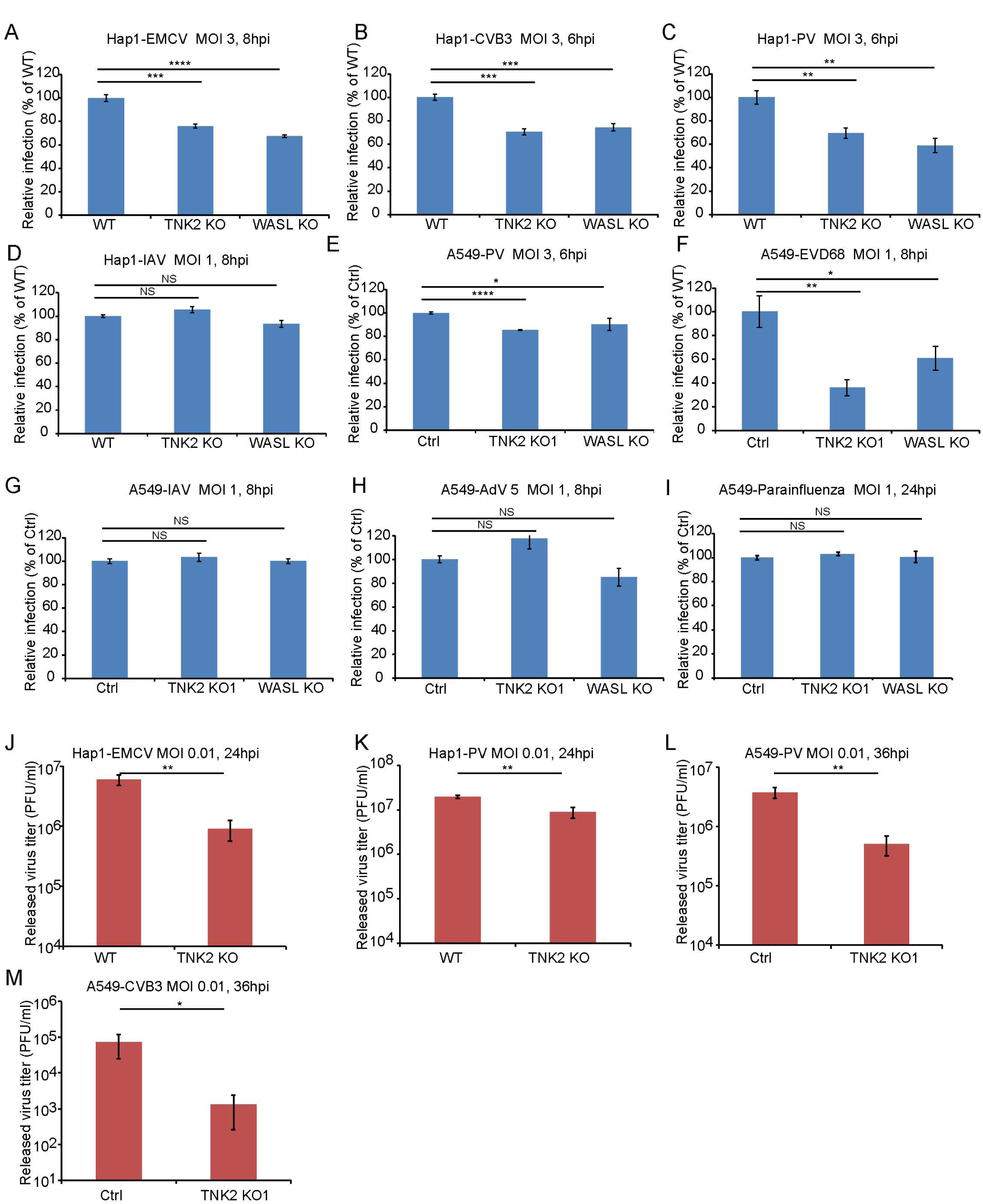
TNK2 and WASL are critical for multiple picornavirus infection on both A5 4 9 and Hap1 cell. (A-D) FACS quantification of EMCV, GVB3, Poliovirus (PV), and influenza A virus (IAV) virus infection on WT (wild type), TNK2 KO and WASL KO Hap1 cells. (E-I) FACS quantification of poliovirus (PV), enterovirus D68 (EVD68), influenza A virus (IAV) virus, Adenovirus type 5 (AdV5) and parainfluenza virus infection on Ctrl, TNK2 KO and WASL KO A5 4 9 cells. (J) EMCV growth titration on WT and TNK2 KO Hap1 cells at 24 hours post infection. (K) Poliovirus growth titration on WT and TNK2 KO Hap1 cells at 24 hours post infection. (L) Poliovirus growth titration on Ctrl and TNK2 KO1 A5 4 9 cells at 36 hours post infection. (M) CVB3 growth titration on Ctrl and TNK2 KO1 A5 4 9 cells at 36 hours post infection. (A-L) Error bars represent standard deviation of three replicates. The data shown is representative of at least two independent experiments. *: P<0.05, **: P<0.01, ***: P<0.001, ****: P<0.0001, NS: not significant (P>0.05).

**Figure 1-figure supplement 3.**
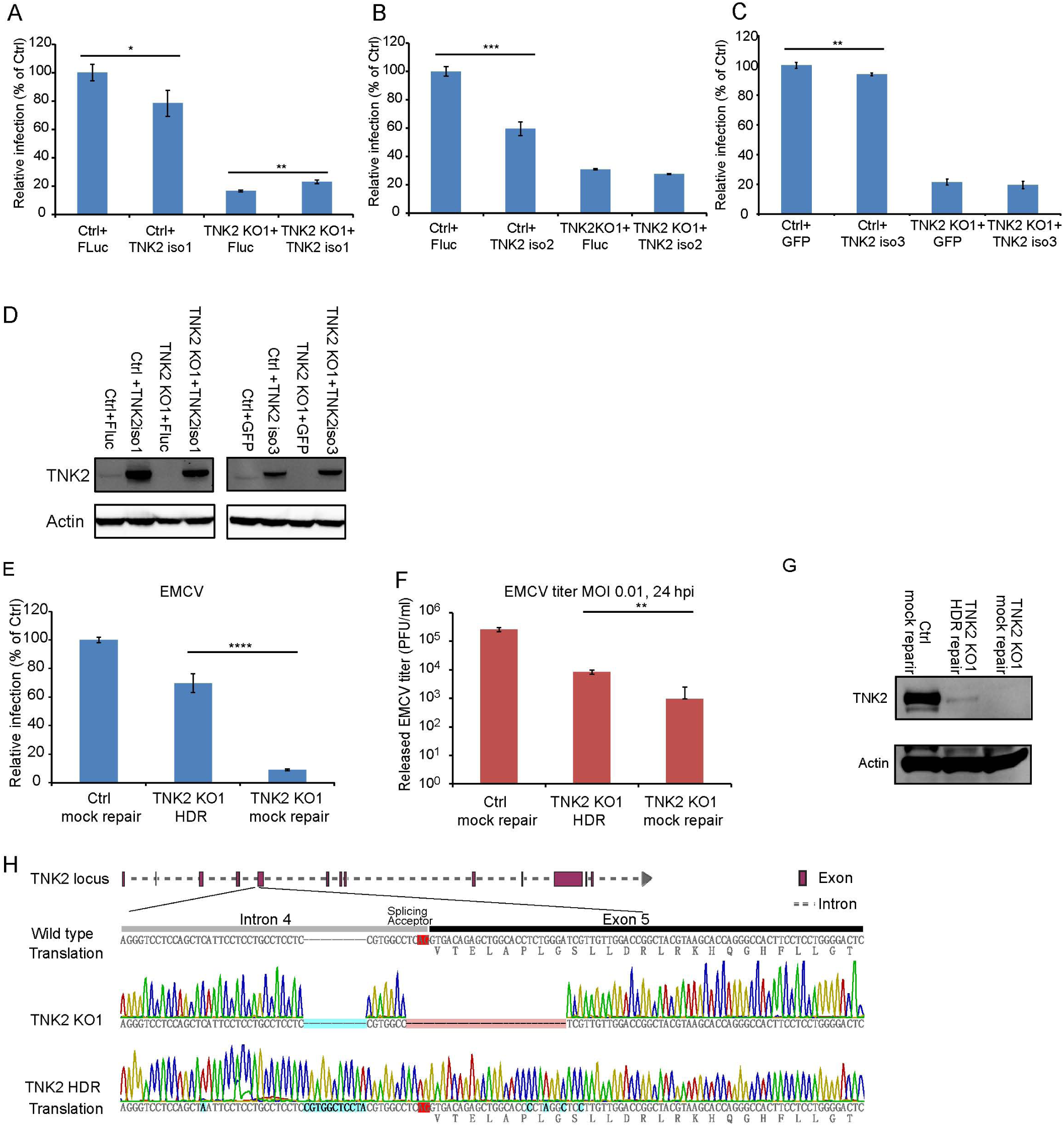
EMCV virus infection of TNK2 rescue on knockout and control cells. (A-C) Quantification of EMCV infection on TNK2 KO1 and Ctrl cells transduced with TNK2 isoform 1, 2 and 3 and Fluc. Cells were infected with EMCV at an MOI of 1 and quantified at 10 hours post infection. (D) Expression of TNK2 in Ctrl and TNK2 KO1 cells transduced with lentivirus expressing TNK2 isoform 1, isoform 3, and Flue or GFP by Western blot. (E) FACS quantification of EMCV positive cells for Ctrl mock repair, TNK2 KO1 HDR, and TNK2 KO1 mock repair cells 10 hours post infection at an MOI of 1. Mock repair: cells subjected to the same HDR genome editing but with non-specific targeting sgRNA. (F) EMCV growth titration on Ctrl mock repair, TNK2 KO1 HDR (homologous template direcfed recombination), and TNK2 KO1 mock repair cells at 24 hours post infection. (G) TNK2 protein expression in Ctrl mock repair, TNK2 KO1 HDR cells and TNK2 KO1 mock repair cells. Cells lysates were analyzed by Western blot. (H) Sequence alignment of TNK2 KO1 cells and TNK2 KO1 HDR (homologous template directed recombination) repaired cells. Splice acceptor is marked with red; synonymous mutations introduced by design and random insertion mutations are marked with blue. (A-C, E, F) Error bars represent standard deviation of three replicates. The data shown are representative of at least two independent experiments. Flue: firefly luciferase. *: P<0.05, **: P<0.01, ***: P<0.001, NS: not significant (P>0.05).

**Figure 2-figure supplement.**
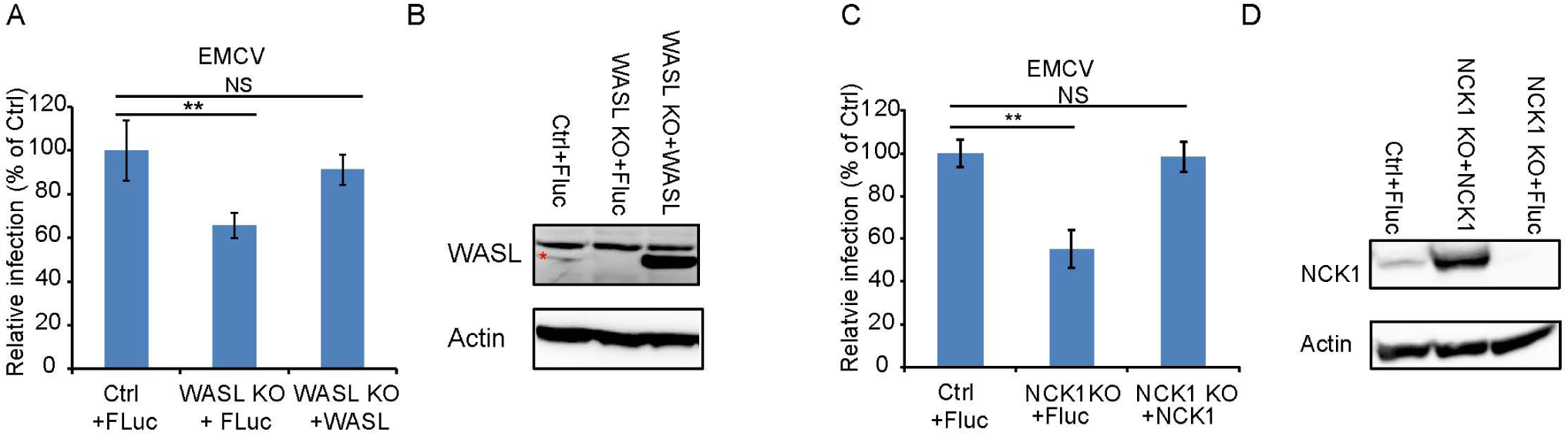
EMCV infection of WASL and NCK1 rescued knock out cells (A) FACS quantification of EMCV positive cells for lentivirus-mediated WASL rescue in WASL KO and Ctrl cells at an MOI of 1. Flue: firefly luciferase. (B) Western blot detection of lentivirus-mediated WASL expression in WASL KO and Ctrl cells. (C) FACS quantification of EMCV positive cells for lentivirus-mediated NCK1 rescue in NCK1 KO and Ctrl cells at an MOI of 1. (D) Western blot detection of lentivirus mediated NCK1 expression in NCK1 KO and Ctrl cells. Error bars represent standard deviation of three replicates. The data shown is representatives of at least two independent experiments. **: P<0.01, NS: not significant (P>0.05).

**Figure 3-figure supplement1.**
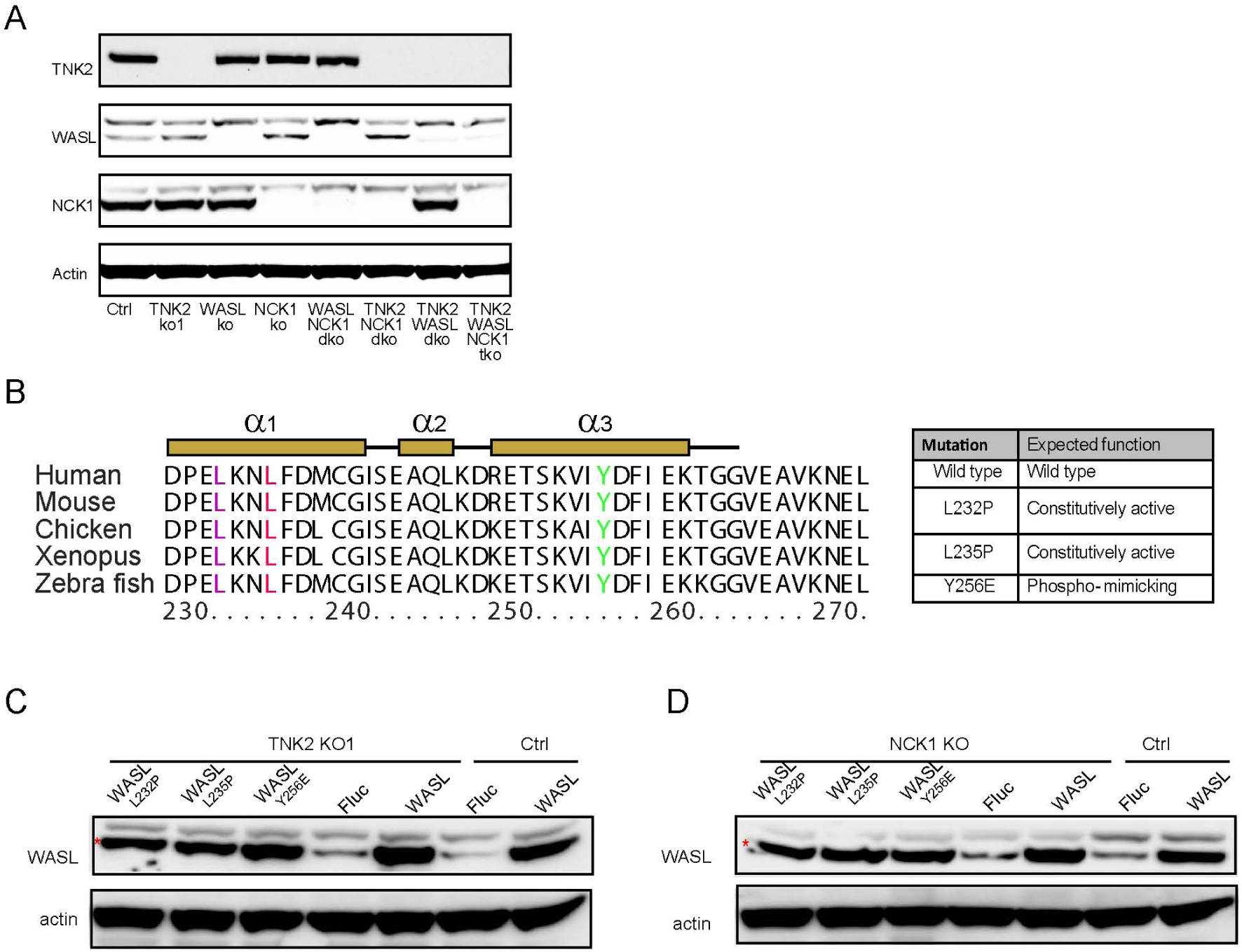
Gene expression in knock out cells and constitutively active WASL expression. (A) Western blot detection ofTNK2, WASL, and NCK1 expression on single, double, triple gene knockout and Ctrl cells. (B) Sequence alignment ofWASL protein sequences from different vertebrate species. Table indicates specific mutation that have constitutive activity. (C) Expression of WASL in Ctrl and TNK2 KO1 cells transduced with lentivirus expressing constitutively active WASL, wild type WASL, or Fluc. Cell lysates were analyzed by Western blot. (D) Expression of WASL in Ctrl and NCK1 KO cells transduced with lentivirus expressing constitutively active WASL, wild type WASL, or Fluc. Cell lysates were analyzed by Western blot. (C, D) The red asterisks indicate WASL protein band.

**Figure 3-figure supplement3.**
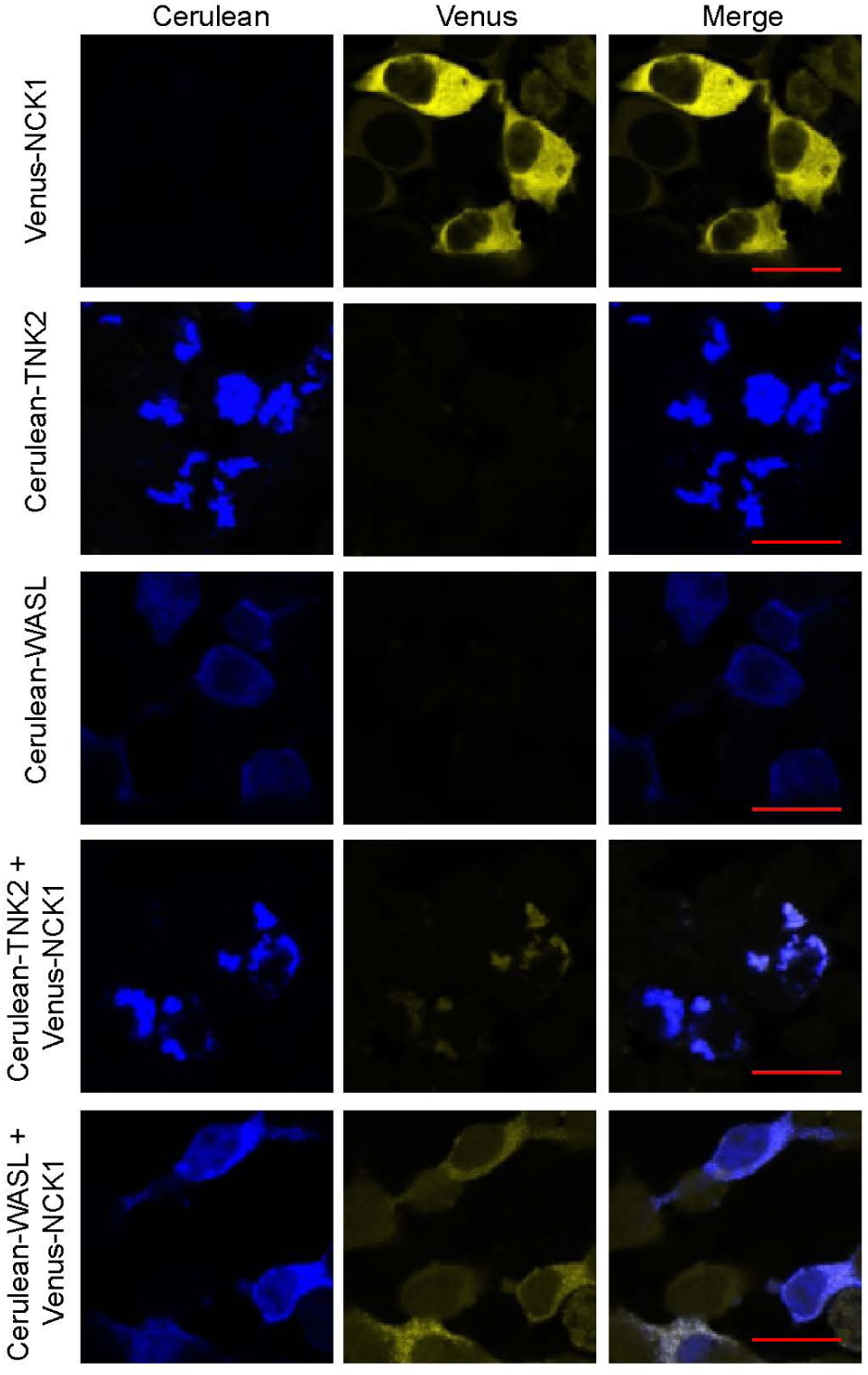
Co-localization of fluorescently tagged NCK1, TNK2, and WASL expressed in 293T cells by confocal imaging. 293T cells were transfected with Venus-NCK1, Cerulean-TNK2, Cerulean-WASL individually, Venus-NCK1 with Cerulean-TNK2 or Venus-NCK1 with Cerulean-WASL. Cells were imaged by confocal 24 hours after transfection. Scale bars represent 20 µm.

**Figure 3-figure supplement 3.**
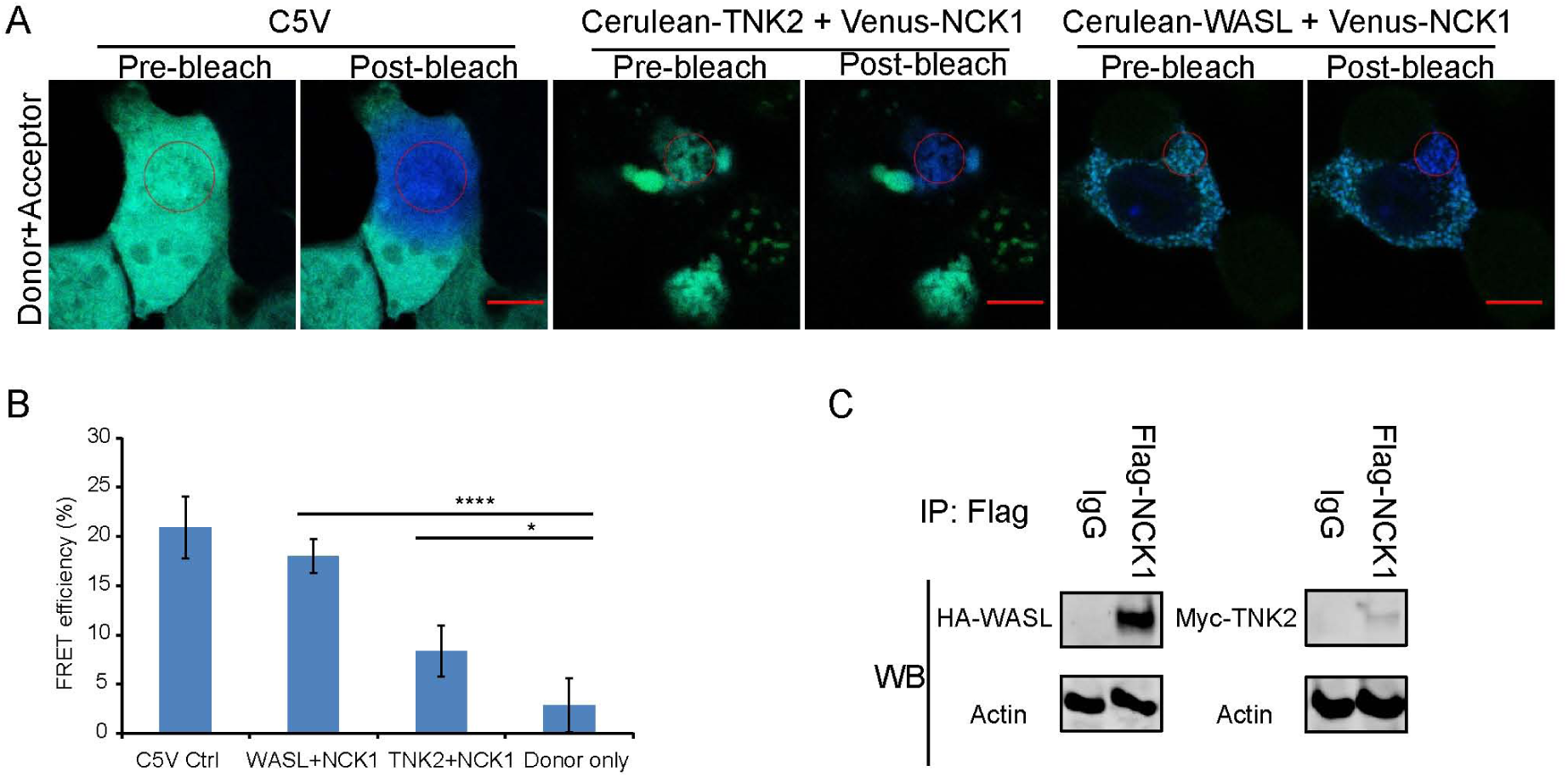
TNK2 and WASL directly interact with NCK1. (A) FRET imaging of C5V positive control, NCK1 and TNK2, and NCK1 and WASL expression in 293T cells before and after acceptor photo bleach. The red circles indicate photo bleached areas. Scale bars represents 10 µm. (B) Quantification of FRET efficiency of C5V positive control, NCK1 and TNK2, and NCK1 and WASL expression in 293T cells. Error bars for FRET efficiency represent standard deviation of average from four individually bleached images. *: P<0.05, ****: P<0.0001, NS: not significant (P>0.05). (C) lmmunoprecipitation of FLAG-tagged NCK1 with HA-tagged WASL and FLAG-tagged NCK1 with Myc-tagged TNK2.

**Figure 4-figure supplement.**
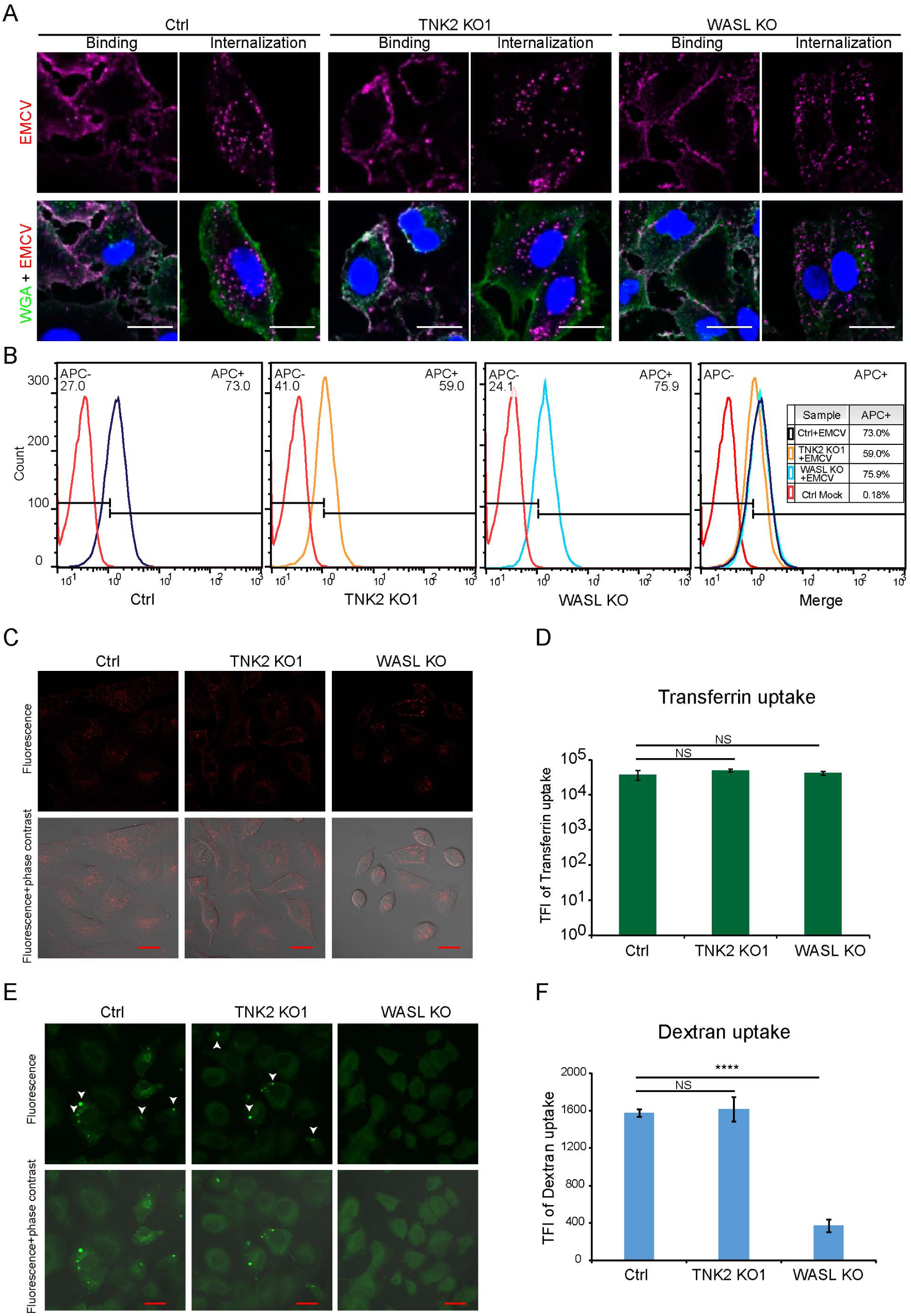
EMCV internalization, transferrin, and dextran uptake in Ctrl, TNK2 KO1 and WASL KO cells. (A) Labeled EMCV viruses bind and internalize in TNK2 KO1, WASL KO and Ctrl cells. Scale bars represents 20 µm. (B) FACS plot of labeled EMCV internalization in TNK2 KO1, WASL KO, and Ctrl cells. (C) Fluorescent images of transferrin uptake in Ctrl, TNK2 KO1, and WASL KO cells. Scale bars represent 20 µm. (D) Quantification of transferrin uptake by flow cytometry. Error bars represent standard deviation of three replicates. The data shown is representative of two independent experiments. TFI: total fluorescence intensity. NS: not significant (P>0.05). (E) Fluorescent images of dextran uptake in Ctrl, TNK2 KO1, and WASL KO cells. White arrows indicate macropinosomes. Scale bars represent 20 µm. (F) Quantification of Dextran uptake by flow cytometry. Error bars represent standard deviation of three replicates. The data shown is representative of two independent experiments. TFI: total fluorescence intensity. NS: not significant (P>0.05), ****: P<0.0001.

**Figure 5-figure supplement.**
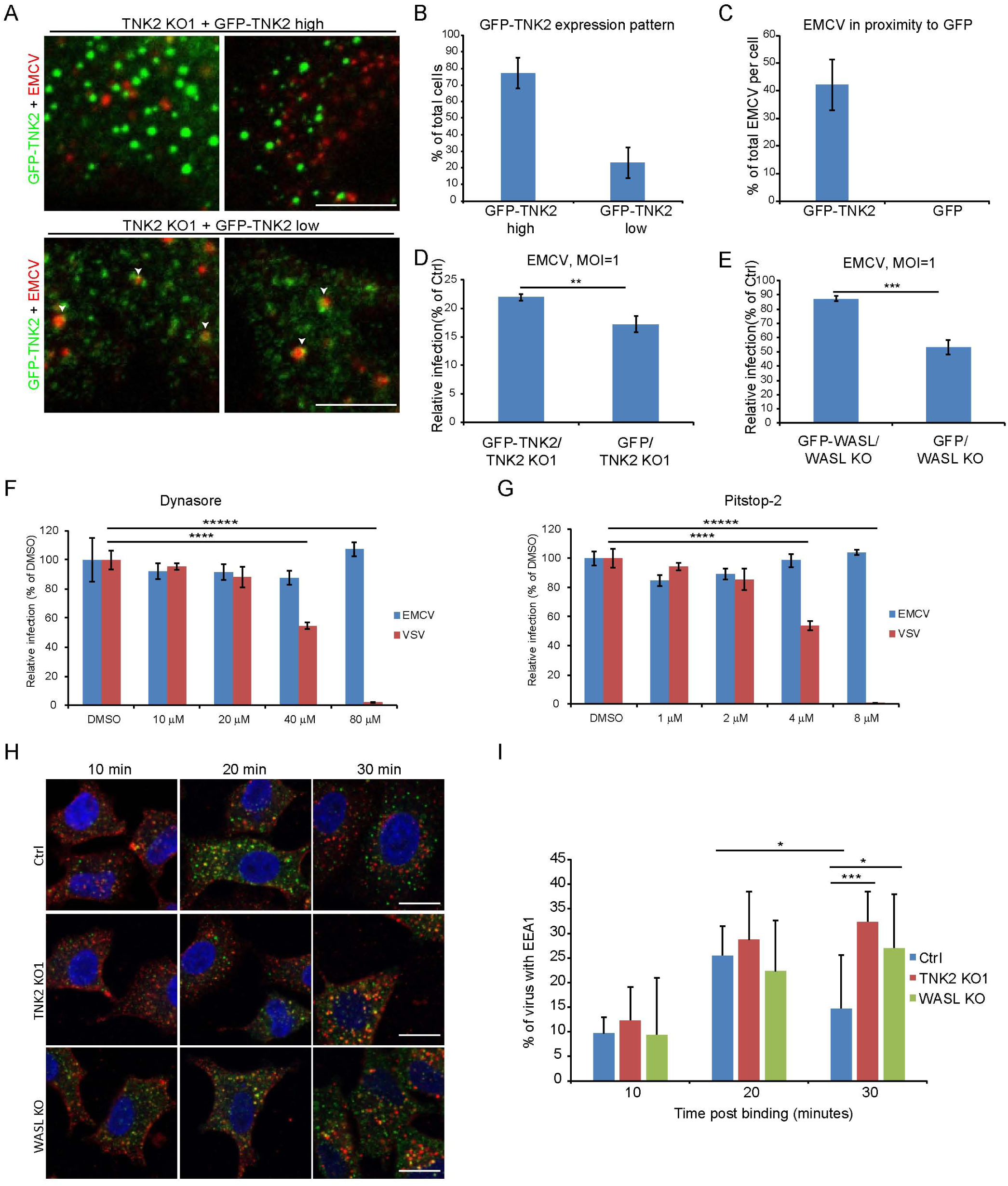
Localization of GFP-TNK2 with fluorescently labeled EMCV. (A) Fluorescent images of different GFP-TNK2 expression patterns in TNK2 KO1 cells and its localization with fluorescently labeled EMCV. White arrow heads indicate EMCVparticles in proximity to GFP-TNK2. Scale bars represent 10µm. (B) Quantification of GFP-TNK2 high and low expression patterns in TNK2 KO1 cells. (C) Quantification of EMCV particles in proximity to GFP-TNK2 in TNK2 KO1 cells express low GFP-TNK2 compared to EMCV particles in proximity to GFP in TNK2 KO1 cells express GFP. (D) Quantification of EMCV infection on TNK2 KO1 cells transduced with lentivirus expressing GFP-TNK2 or GFP at 10hours post infection. Error bars represent standard deviation of three replicates. The data shown is representative of two independent experiments. **:P<0.01. (E) Quantification of EMCV infection on WASL KO cells transduced with lentivirus expressing GFP-WASL or GFP at 10 hours post infection. Error bars represent standard deviation of three replicates. The data shown is representative of two independent experiments. ***: P<0.001. (F) Dynasore inhibition of EMCV and VSV infection on naïve A549 cells. (G) Pitstop-2 inhibition of EMCV and VSV infection on naïve A549 cells. (H) Fluorescent images of EEA1 co-localization with fluorescently labeled EMCV at 10, 20, and 30 minutes post internalization. Scale bars represent 20 µm. (I) Quantification of EEA1 co-localization with fluorescently labeled EMCV at 10, 20, and 30 minutes post internalization. Percentage of EMCV co-localized with EEA1 were quantified by image analysis. Error bars represent standard deviation of three experiments. *:P<0.05, ***: P<0.001.

## References

1. Melnick JL (1983) Portraits of viruses: the picornaviruses. Intervirology 20(2-3):61–100.

2. Yamauchi Y & Helenius A (2013) Virus entry at a glance. J Cell Sci 126(Pt 6):1289–1295.

3. Madshus IH, Olsnes S, & Sandvig K (1984) Requirements for entry of poliovirus RNA into cells at low pH. The EMBO journal 3(9):1945–1950.

4. Brandenburg B, et al. (2007) Imaging poliovirus entry in live cells. PLoS biology 5(7):e183.

5. Bergelson JM & Coyne CB (2013) Picornavirus entry. Advances in experimental medicine and biology 790:24–41.

6. Coyne CB & Bergelson JM (2006) Virus-induced Abl and Fyn kinase signals permit coxsackievirus entry through epithelial tight junctions. Cell 124(1):119–131.

7. Coyne CB, Shen L, Turner JR, & Bergelson JM (2007) Coxsackievirus entry across epithelial tight junctions requires occludin and the small GTPases Rab34 and Rab5. Cell host & microbe 2(3):181–192.

8. Huber SA (1994) VCAM-1 is a receptor for encephalomyocarditis virus on murine vascular endothelial cells. Journal of virology 68(6):3453–3458.

9. Madshus IH, Olsnes S, & Sandvig K (1984) Different pH requirements for entry of the two picornaviruses, human rhinovirus 2 and murine encephalomyocarditis virus. Virology 139(2):346–357.

10. Jiang H, Chen K, Sandoval LE, Leung C, & Wang D (2017) An Evolutionarily Conserved Pathway Essential for Orsay Virus Infection of Caenorhabditis elegans. mBio 8(5).

11. Galisteo ML, Yang Y, Urena J, & Schlessinger J (2006) Activation of the nonreceptor protein tyrosine kinase Ack by multiple extracellular stimuli. Proceedings of the National Academy of Sciences of the United States of America 103(26):9796–9801.

12. Donnelly SK, Weisswange I, Zettl M, & Way M (2013) WIP provides an essential link between Nck and N-WASP during Arp2/3-dependent actin polymerization. Current biology: CB 23(11):999–1006.

13. Liu Y, et al. (2010) Dasatinib inhibits site-specific tyrosine phosphorylation of androgen receptor by Ack1 and Src kinases. Oncogene 29(22):3208–3216.

14. Mahajan NP, Whang YE, Mohler JL, & Earp HS (2005) Activated tyrosine kinase Ack1 promotes prostate tumorigenesis: role of Ack1 in polyubiquitination of tumor suppressor Wwox. Cancer Res 65(22):10514–10523.

15. Yokoyama N, Lougheed J, & Miller WT (2005) Phosphorylation of WASP by the Cdc42-associated kinase ACK1: dual hydroxyamino acid specificity in a tyrosine kinase. The Journal of biological chemistry 280(51):42219–42226.

16. Fujimoto Y, et al. (2011) A single nucleotide polymorphism in activated Cdc42 associated tyrosine kinase 1 influences the interferon therapy in hepatitis C patients. Journal of hepatology 54(4):629–639.

17. Konig R, et al. (2010) Human host factors required for influenza virus replication. Nature 463(7282):813–817.

18. Karlas A, et al. (2010) Genome-wide RNAi screen identifies human host factors crucial for influenza virus replication. Nature 463(7282):818–822.

19. Lupberger J, et al. (2011) EGFR and EphA2 are host factors for hepatitis C virus entry and possible targets for antiviral therapy. Nature medicine 17(5):589–595.

20. Massaad MJ, Ramesh N, & Geha RS (2013) Wiskott-Aldrich syndrome: a comprehensive review. Annals of the New York Academy of Sciences 1285:26–43.

21. Teo M, Tan L, Lim L, & Manser E (2001) The tyrosine kinase ACK1 associates with clathrin-coated vesicles through a binding motif shared by arrestin and other adaptors. The Journal of biological chemistry 276(21):18392–18398.

22. Frischknecht F, et al. (1999) Actin-based motility of vaccinia virus mimics receptor tyrosine kinase signalling. Nature 401(6756):926–929.

23. Dodding MP & Way M (2009) Nck- and N-WASP-dependent actin-based motility is conserved in divergent vertebrate poxviruses. Cell host & microbe 6(6):536–550.

24. Oppliger J, Torriani G, Herrador A, & Kunz S (2016) Lassa Virus Cell Entry via Dystroglycan Involves an Unusual Pathway of Macropinocytosis. Journal of virology 90(14):6412–6429.

25. Wolf YI, et al. (2018) Origins and Evolution of the Global RNA Virome. mBio 9(6).

26. Zerbino DR, et al. (2018) Ensembl 2018. Nucleic acids research 46(D1):D754–D761.

27. Mahajan K, et al. (2010) Effect of Ack1 tyrosine kinase inhibitor on ligand-independent androgen receptor activity. The Prostate 70(12):1274–1285.

28. Peterson JR, et al. (2004) Chemical inhibition of N-WASP by stabilization of a native autoinhibited conformation. Nature structural & molecular biology 11(8):747–755.

29. Yokoyama N & Miller WT (2003) Biochemical properties of the Cdc42-associated tyrosine kinase ACK1. Substrate specificity, authphosphorylation, and interaction with Hck. J Biol Chem 278(48):47713–47723.

30. Adamovich DA, et al. (2009) Activating mutations of N-WASP alter Shigella pathogenesis. Biochemical and biophysical research communications 384(3):284–289.

31. Keszei M, et al. (2018) Constitutive activation of WASp in X-linked neutropenia renders neutrophils hyperactive. The Journal of clinical investigation 128(9):4115–4131.

32. Rohatgi R, Nollau P, Ho HY, Kirschner MW, & Mayer BJ (2001) Nck and phosphatidylinositol 4,5-bisphosphate synergistically activate actin polymerization through the N-WASP-Arp2/3 pathway. The Journal of biological chemistry 276(28):26448–26452.

33. Shen H, et al. (2011) Constitutive activated Cdc42-associated kinase (Ack) phosphorylation at arrested endocytic clathrin-coated pits of cells that lack dynamin. Molecular biology of the cell 22(4):493–502.

34. Berry GE & Tse LV (2017) Virus Binding and Internalization Assay for Adeno-associated Virus. Bio Protoc 7(2).

35. Hackett BA, et al. (2015) RNASEK is required for internalization of diverse acid-dependent viruses. Proceedings of the National Academy of Sciences of the United States of America 112(25):7797–7802.

36. Grovdal LM, Johannessen LE, Rodland MS, Madshus IH, & Stang E (2008) Dysregulation of Ack1 inhibits down-regulation of the EGF receptor. Experimental cell research 314(6):1292–1300.

37. Innocenti M, et al. (2005) Abi1 regulates the activity of N-WASP and WAVE in distinct actin-based processes. Nature cell biology 7(10):969–976.

38. Fernandez-Puentes C & Carrasco L (1980) Viral infection permeabilizes mammalian cells to protein toxins. Cell 20(3):769–775.

39. Porotto M, Palmer SG, Palermo LM, & Moscona A (2012) Mechanism of fusion triggering by human parainfluenza virus type III: communication between viral glycoproteins during entry. The Journal of biological chemistry 287(1):778–793.

40. Staring J, et al. (2017) PLA2G16 represents a switch between entry and clearance of Picornaviridae. Nature 541(7637):412–416.

41. Glaser W & Skern T (2000) Extremely efficient cleavage of eIF4G by picornaviral proteinases L and 2A in vitro. FEBS Lett 480(2-3):151–155.

42. Carthy CM, et al. (1998) Caspase activation and specific cleavage of substrates after coxsackievirus B3-induced cytopathic effect in HeLa cells. Journal of virology 72(9):7669–7675.

43. Jones S, Cunningham DL, Rappoport JZ, & Heath JK (2014) The non-receptor tyrosine kinase Ack1 regulates the fate of activated EGFR by inducing trafficking to the p62/NBR1 pre-autophagosome. J Cell Sci 127(Pt 5):994–1006.

44. Prieto-Echague V, Gucwa A, Brown DA, & Miller WT (2010) Regulation of Ack1 localization and activity by the amino-terminal SAM domain. BMC Biochem 11:42.

45. Howlin J, Rosenkvist J, & Andersson T (2008) TNK2 preserves epidermal growth factor receptor expression on the cell surface and enhances migration and invasion of human breast cancer cells. Breast cancer research: BCR 10(2):R36.

46. Mahajan NP, et al. (2007) Activated Cdc42-associated kinase Ack1 promotes prostate cancer progression via androgen receptor tyrosine phosphorylation. Proceedings of the National Academy of Sciences of the United States of America 104(20):8438–8443.

47. Rohatgi R, Ho HY, & Kirschner MW (2000) Mechanism of N-WASP activation by CDC42 and phosphatidylinositol 4, 5-bisphosphate. The Journal of cell biology 150(6):1299–1310.

48. Galletta BJ, Chuang DY, & Cooper JA (2008) Distinct roles for Arp2/3 regulators in actin assembly and endocytosis. PLoS biology 6(1):e1.

49. Stradal TE, et al. (2004) Regulation of actin dynamics by WASP and WAVE family proteins. Trends in cell biology 14(6):303–311.

50. Prieto-Echague V & Miller WT (2011) Regulation of ack-family nonreceptor tyrosine kinases. Journal of signal transduction 2011:742372.

51. Jose AM, Kim YA, Leal-Ekman S, & Hunter CP (2012) Conserved tyrosine kinase promotes the import of silencing RNA into Caenorhabditis elegans cells. Proceedings of the National Academy of Sciences of the United States of America 109(36):14520–14525.

52. Bladt F, et al. (2003) The murine Nck SH2/SH3 adaptors are important for the development of mesoderm-derived embryonic structures and for regulating the cellular actin network. Molecular and cellular biology 23(13):4586–4597.

53. Kim HS, et al. (2017) CRISPR/Cas9-mediated gene knockout screens and target identification via whole-genome sequencing uncover host genes required for picornavirus infection. The Journal of biological chemistry 292(25):10664–10671.

54. Bazzone LE, et al. (2019) A Disintegrin and Metalloproteinase 9 Domain (ADAM9) Is a Major Susceptibility Factor in the Early Stages of Encephalomyocarditis Virus Infection. mBio 10(1).

55. Winston WM, Molodowitch C, & Hunter CP (2002) Systemic RNAi in C. elegans requires the putative transmembrane protein SID-1. Science 295(5564):2456–2459.

56. Nguyen TA, et al. (2017) SIDT2 Transports Extracellular dsRNA into the Cytoplasm for Innate Immune Recognition. Immunity 47(3):498–509 e496.

57. Franz CJ, et al. (2014) Orsay, Santeuil and Le Blanc viruses primarily infect intestinal cells in Caenorhabditis nematodes. Virology 448:255–264.

58. Edelman BL & Redente EF (2018) Isolation and Characterization of Mouse Fibroblasts. Methods Mol Biol 1809:59–67.

59. Jacobi AM, et al. (2017) Simplified CRISPR tools for efficient genome editing and streamlined protocols for their delivery into mammalian cells and mouse zygotes. Methods 121-122:16–28.

60. Wang P, et al. (2015) Nlrp6 regulates intestinal antiviral innate immunity. Science 350(6262):826–830.

